# Humidity determines penetrance of a latitudinal gradient in genetic selection on the microbiota by *Drosophila melanogaster*

**DOI:** 10.1101/2024.05.02.591907

**Authors:** Caroline Massey, Maggie E. Nosker, Joseph Gale, Shayna Scott, Carson J. Walker, Aubrey Cluff, Susan Wilcox, Amanda Morrison, Sarah J. Gottfredson Morgan, Jack Beltz, Paul Schmidt, John M. Chaston

## Abstract

The fruit fly *Drosophila melanogaster* is a model for understanding how hosts and their microbial partners interact as the host adapts to wild environments. These interactions are readily interrogated because of the low taxonomic and numeric complexity of the flies’ bacterial communities. Previous work has established that host genotype, the environment, diet, and interspecies microbial interactions can all influence host fitness and microbiota composition, but the specific processes and characters mediating these processes are incompletely understood. Here, we compared the variation in microbiota composition between wild-derived fly populations when flies could choose between the microorganisms in their diets and when flies were reared under environmental perturbation (different humidities). We also compared the colonization of the resident and transient microorganisms. We show that the ability to choose between microorganisms in the diet and the environmental condition of the flies can influence the relative abundance of the microbiota. There were also key differences in the abundances of the resident and transient microbiota. However, the microbiota only differed between populations when the flies were reared at humidities at or above 50% relative humidity. We also show that elevated humidity determined the penetrance of a gradient in host genetic selection on the microbiota that is associated with the latitude the flies were collected from. Finally, we show that the treatment-dependent variation in microbiota composition is associated with variation in host stress survival. Together, these findings emphasize that host genetic selection on the microbiota composition of a model animal host can be patterned with the source geography, and that such variation has the potential to influence their survival in the wild.

**Importance:** The fruit fly *Drosophila melanogaster* is a model for understanding how hosts and their microbial partners interact as hosts adapt in wild environments. Our understanding of what causes geographic variation in the fruit fly microbiota remains incomplete. Previous work has shown that the *D. melanogaster* microbiota has relatively low numerical and taxonomic complexity. Variation in the fly microbiota composition can be attributed to environmental characters and host genetic variation, and variation in microbiota composition can be patterned with the source location of the flies. In this work we explored three possible causes of patterned variation in microbiota composition. We show that host feeding choices, the host niche colonized by the bacteria, and a single environmental character can all contribute to variation in microbiota composition. We also show that penetrance of latitudinally-patterned host genetic selection is only observed at elevated humidities. Together, these results identify several factors that influence microbiota composition in wild fly genotypes and emphasize the interplay between environmental and host genetic factors in determining the microbiota composition of these model hosts.

## Introduction

The microbiota is the community of microorganisms associated with an abiotic or biotic (e.g. host) environment. Host interactions with the microbiota have been studied in diverse animal species that have relevance to human health or are associated with various host conditions and diseases (1–8). For example, host-associated microbial communities play a crucial role in shaping host health and behavior in the fruit fly *Drosophila melanogaster*, including life history traits such as lifespan and development rate (9–16). As in many animals, fruit fly microbiota composition is not strictly programmed from birth but is flexible and influenced by various conditions experienced by the flies, including the host’s environment and biology (9, 17–21). Conversely, environment-dependent variation in microbiota composition can influence the host’s traits (10). To better understand and predict host responses to variation in microbiota composition we investigated how host genetic selection, bacterial colonization proficiency, and environmental conditions, impact microbiota composition in wild flies from different locations across a latitudinal gradient in the eastern United States (USA).

In the fruit fly, *D. melanogaster,* the microbiota composition is both relatively low in diversity and readily manipulated. (22–26). Their microbiota composition is frequently dominated by acetic acid bacteria (AAB), lactic acid bacteria (LAB), and enterobacteria (27–32). Some microorganisms colonize the flies only transiently, for no longer than the bulk flow of diet through the fly gastrointestinal tract, but many members of the microbiota can colonize the flies for much longer (20, 21, 33, 34). Most colonizing bacteria are spatially patterned to specific niches within distinct host foregut tissues (35–38). Resident microorganisms can readily be distinguished by measuring bacterial loads in flies after a brief starvation period during which transient microorganisms are defecated. Most microbiota members can be readily cultured in the laboratory separately from their animal host, and the host can be reared with defined communities of these microorganisms by inoculating bleached fly embryos with suspensions of one or more bacterial species (39). Together, these characteristics make the partnership highly amenable to laboratory study.

Previous work has shown that host genotype, environmental characters, diet, and microbe-microbe interactions are all factors that can contribute to variation in the *D. melanogaster* microbiota composition. For example, temperature influences shifts in the microbial communities of *Drosophila* species (18, 19) and some fly genes have been identified that influence microbiota composition (9). Fly feeding preferences for different microbes in their diets provides a possible explanation for how fly genotype may select on the microbiota composition (40). Diet and inter-species microbe-microbe interactions also play key roles in determining of microbiota composition (11, 25, 27, 36, 41–48). Together, these previous efforts have helped to explain some factors that determine the microbiota composition of adult flies.

Despite the progress cited above, our understanding of what causes variation in the microbiota of wild *D. melanogaster* is incompletely understood. A 16S sequencing analysis of the microbiota of six wild fly populations in the eastern USA showed a gradient in fly microbiota composition where the AAB were more abundant in flies at low latitudes and the LAB were more abundant in flies at high latitudes (10). We previously suggested that at least some of this variation was due to host genetic selection on the microbiota because fly lines from Maine and Florida, USA, reared from the embryonic stage with the same starting microorganisms, had different microbiota composition as adults (10). The low-latitude population from Florida had a higher relative abundance of AAB, whereas the high-latitude population from Maine had more LAB (10). However, this latitudinal pattern is not universal since a more recent analysis of microbiota composition in wild flies on two diets across many of the same locations revealed almost no LAB and no pattern in the AAB:LAB ratio (32). Additionally, the gnotobiotic fly analysis we conducted previously used only two genotypes and was therefore unable to test if there is gradient in host genetic selection on the microbiota. Therefore, there are key gaps in our understanding of what determines the microbiota composition of flies in the wild.

In this study, we identify humidity as a major factor that influences how *D. melanogaster* in the eastern USA genetically select for their microbiota composition. We collected wild flies from eight locations, creating common garden populations of the flies from each location, and compared their microbiota composition under three conditions: 1) when the flies could or could not choose between microorganisms in their diet; 2) when the transient and colonizing microorganisms were distinguished from each other; and 3) when the flies were temporarily reared at different humidities. In each condition we tested if the condition influenced the microbiota composition and if the populations had different microbiota composition from each other. We determined that microbiota composition varied in response to fly choice and humidity treatments and that at high humidities the transient, but not the colonizing, portion of the microbiota covaried significantly with the source latitude of the fly populations. These findings reinforce that in *D. melanogaster* host genotype can select the microbiota in a way that is patterned with the animal’s geographic location.

## Materials and Methods

### General Rearing and Culture Conditions

For general conditions, flies were reared at 25°C in ambient humidity (∼25% relative humidity (RH)) on a 12-hour light: 12-hour dark cycle. A *Wolbachia*-free laboratory CantonS fly line originally obtained from Mariana Wolfner was reared on a yeast-glucose (Y-G) diet (10% Brewer’s yeast (MP Biomedicals 903312), 10% glucose (Sigma, 102506465), 1% agar (FlyStuff 66-103)) with acid preservative added at 0.42% propanoic acid and 0.04% phosphoric acid. Wild collections of flies were reared on a molasses diet of 7.05% molasses (FlyStuff 62-117), 5.48% cornmeal (FlyStuff 62-100), 1.31% Brewer’s yeast, 0.74% soy flour (FlyStuff 62-115), 0.43% agar, 0.18% Tegosept (Apex 20-258), and 2.91% 190 proof ethanol.

In this study we used bacterial strains *Acetobacter tropicalis* DmCS_006*, Acetobacter sp.* DsW_54*, Acetobacter sp.* DmW_125*, A. orientalis DmW_045, Lactiplantibacillus plantarum* DmCS_001*, Weissella paramesenteroides* DmW_115*, and Leuconostoc suionicum* DmW_098. Each was cultured on modified de Man-Rogosa-Sharpe (mMRS) medium (Criterion C5932) for 2-3 days at 30°C. Solid medium was supplemented with 1.5% agar before autoclaving. The *Acetobacter* strains were grown under oxic conditions overnight, with continuous shaking in liquid media and at ambient atmosphere for solid media. The LAB strains were grown under microoxic conditions, with static growth for 1-3 days in liquid media or in an airtight container flooded with CO_2_ for solid media.

To measure the influence of humidity on the *D. melanogaster* microbiota, flies were reared at variable humidities as adults. First, flies were reared from eggs at 25°C, 25% RH, and 12-hour light/dark cycle until the adult flies were 3 days post eclosion. Then, the flies were divided into three groups and either left at ambient humidity (∼ 25% RH) or moved to light- and temperature-controlled incubators set to the same temperature and light cycles but to either 50% or 75% RH. In practice, RH set to 75% fluctuated regularly between 60% RH and 75% RH.

### Establishing a collection of wild-derived fly lines

To capture a range of genetic variation, eight *D. melanogaster* populations were established from wild collections of flies at eight orchards in the eastern USA (Table 1, Fig. 1A). At each orchard, flies were collected with empty 35-ml vials, nets, and hand vacuums from fallen fruits and, where available, compost piles. After several hours, flies were briefly anesthetized using FlyNap (173010, Carolina Biological Supply), and individual females were separated into individual vials containing 7.5 ml of molasses diet. Vials were kept at room temperature and maintained at room temperature on molasses diet in perpetuity as isofemale lines thereafter. From the F1 generation in each vial, males were examined by microscopy to discriminate *D. simulans* and *D. melanogaster*. Each line was assigned as a *D. melanogaster* line if the male fly had a distinct genital arch, and all other lines were discarded. The identity of each line assigned as *D. melanogaster* was confirmed using a PCR assay (49). Five flies from each isofemale line were pooled with five flies from each of four other lines (25 flies from 5 isofemale lines total) for a single DNA extraction. DNA was extracted using a Quick-DNA Fecal/Soil Microbe Miniprep Kit (Zymo D6010). From the extracted DNA, part of the cytochrome C oxidase subunit I gene was amplified using Terra™ PCR Direct Polymerase Mix (639270) with the following primers: HCO (5′-GGT-CAA-CAA-ATC-ATA-AAG-ATA-TTG-G-3′) and LCO (5′-TAA-ACT-TCA-CGG-TGA-CCA-AAA-AAT-CA-3′). Then, a restriction fragment length polymorphism (RFLP) analysis was performed on the PCR product by combining 1µL of *MboII* enzyme, 2 μl of 10× reaction buffer, 7 μl of PCR product; incubating at 37 °C for 1 h; and comparing by gel electrophoresis the digestion pattern with a control digestion (H_2_O added instead of *MboII* enzyme) for each product. Flies were assigned as *D. melanogaster* if there were two bands at ∼ 250 and 450 bp, or as *D. simulans* if they had two bands close to 350 bp or two bands at ∼ 250 and 350 bp. All pools of flies had a single signal of a *D. melanogaster*, providing molecular confirmation for each of the morphological assignments. Therefore, all flies used in this study were *D. melanogaster*.

**Table 1.** *D. melanogaster* populations used in this study.

| *Code in figures* | *Location* | *Orchard name* | *Latitude* | *Longitude* | *# of isofemale lines* |
| --- | --- | --- | --- | --- | --- |
| *ME2* | Etna, ME | Conant Orchards | 44.82 | -69.11 | 5 |
| *ME1* | Bowdoin, ME | Rocky Ridge Orchard | 44.03 | -69.94 | 20 |
| *NH* | Lee, NH | Demeritt Hill Farm | 43.09 | -71.05 | 25 |
| *MA* | Harvard, MA | Westward Orchard | 42.51 | -71.57 | 10 |
| *CT* | Middlefield, CT | Lyman Farms | 41.49 | -72.71 | 12 |
| *PA* | Media, PA | Indian Orchards, Linvilla Orchards | 39.88 | -75.41 | 28 |
| *MD* | Churchville, MD | Lohr's Orchard | 39.54 | -76.24 | 14 |
| *VA* | Charlottesville, VA | Carter Mountain Orchard | 38.07 | -78.38 | 5 |

**Figure 1.**
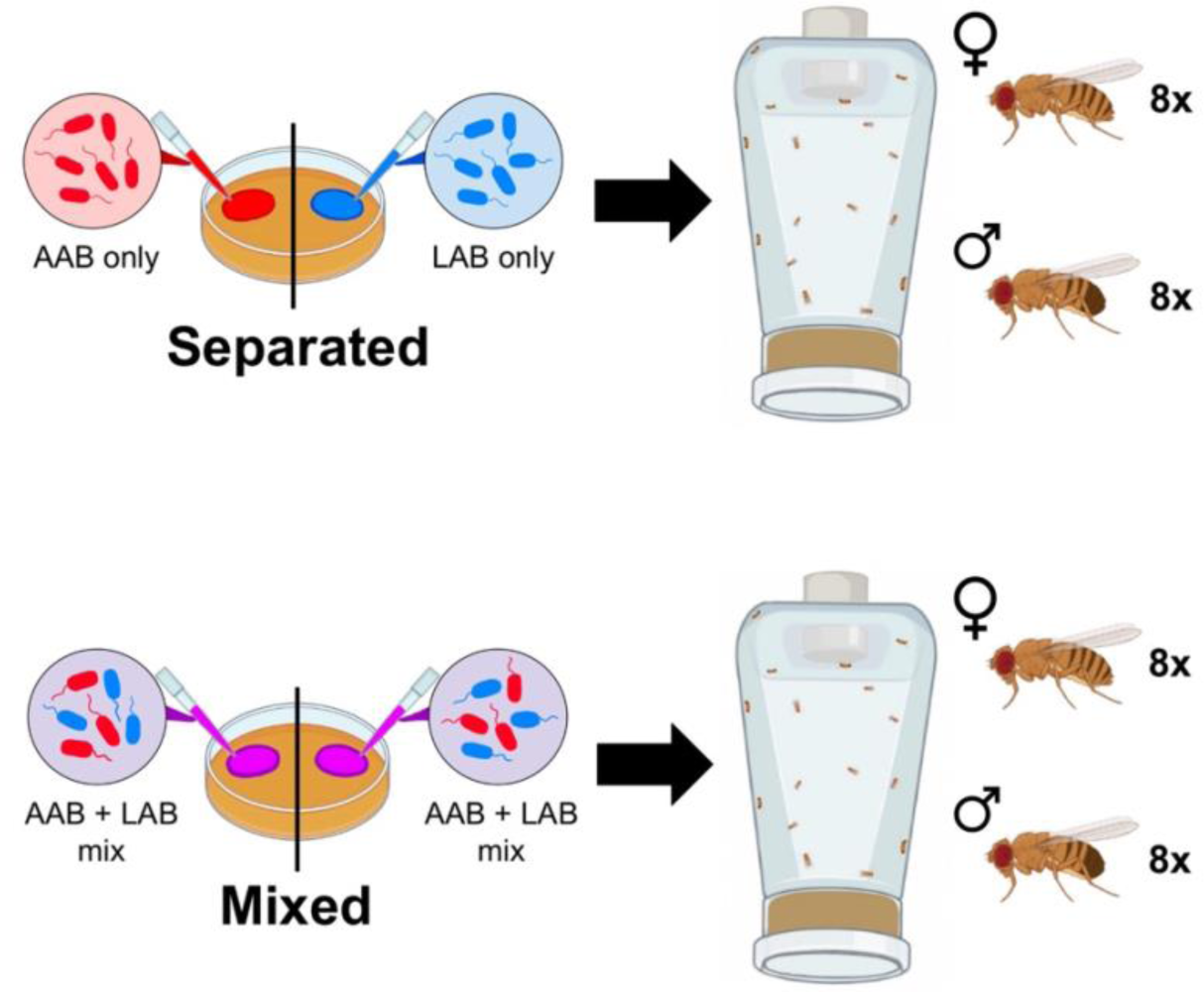
Schematic of bacteria-conditioned diet experiment. Fly genotypes were exposed to diets where the bacterial communities were inoculated separately or mixed together before incoulating to two halve of a diet surface and incubating at 30°C for 24 h. Then, the diets were placed into arenas bearing 16 adults at time = 0. Identically prepared diet plates were replaced after 24 and 48 hours. At 72 hours, adult flies were anesthetized and homogenized to measure their bacterial abundance. Image was created with BioRender.com.

To establish a locale-specific fly population from the individual isofemale lines, we mixed the isofemale lines from each location for 3 generations before performing experiments. From each location, ten young adult (< 7 days old) female and ten young adult male flies from each isofemale line were mixed in a single fly cage on molasses diet (BugDorm, 4S2222). After 1-3 days the diet containing eggs laid by the adults was transferred to a capped 6 oz. fly bottle until the F1 progeny eclosed. Between 2 and 6 days post eclosion, F1 flies were released into a fly cage with molasses diet for 1-3 days to lay eggs, and the diet was again transferred to 6 oz. fly bottles, qualitatively targeting 200-400 eggs per bottle. This process was repeated and either F2 or F3 flies were used to lay the eggs used in ‘conventional’ (no microbiota manipulation) or ‘gnotobiotic’ (reared with a defined microbiota from the embryonic stage) fly experiments.

### Rearing gnotobiotic flies

For gnotobiotic experiments we derived bacteria-free embryos and either inoculated the embryos with specific bacterial strains to create gnotobiotic flies or maintained an axenic (bacteria-free) population without further manipulating the embryos (22). To derive bacteria-free embryos we collected < 18 h old *D. melanogaster* eggs from Y-G-grape juice plates (2 T frozen grape juice concentrate per 500 mL Y-G diet). The eggs were sterilized twice in 0.6% sodium hypochlorite for 150 s each, then rinsed three times with sterile water. Using a paintbrush sterilized in 70% ethanol, 30-50 eggs were transferred to 50 mL conical tubes containing 7.5 mL of autoclave-sterilized molasses diet. Separately, overnight liquid cultures of six bacterial strains (*A. tropicalis* DmCS_006, *Acetobacter* sp. DsW_54, *Acetobacter* sp. DmW_125, *L. plantarum* DmCS_001, *W. paramesenteroides* DmW_115, and *L. suionicum* DmW_098) were normalized to OD_600_ = 0.1, and each normalized microbial suspension was mixed in equivalent ratios, and 50 ml of the mixture was added to the eggs on the diet. In each experiment a few vials were not inoculated with bacteria as a control (axenic) and were allowed mature to adulthood in the absence of bacteria. These were homogenized and dilution plated as described below to confirm fidelity of the sterilization.

### Measuring bacterial abundance

To measure the microbiota composition of adult flies we used a 96-well plate-based approach (36). A pool of two flies was collected to one well in a 96-well plate containing ∼20 ml 0.5 mm glass beads (BioSpec 11079105) and 100 ml phosphate-buffered saline (PBS, 8 g NaCl, 0.2 g KCl, 1.44 g Na_2_HPO_4_, 0.24 g KH_2_PO_4_ L^-1^) on ice. For each treatment we collected two pools of two male flies and two pools of two female flies from each of three replicate vials in each of two or three separate experiments. After flies from all treatments were collected to distinct wells, the plates were heat sealed with a Thermal Bond Heat Sealing Foil (4titude, 4ti-0591) using an Eppendorf HeatSealer S100, homogenized at 1750 rpm for 5 minutes in a SPEX SamplePrep Geno/Grinder 2010, and centrifuged briefly at <4,000 x g to remove the homogenate from the foil lid. Using a 96-well pipettor, plates containing flies were diluted 15.9-fold, followed by two rounds of 8-fold dilutions in 1X phosphate-buffered saline (PBS) while plates containing fly diets were diluted in four sequential 8-fold dilutions to achieve a target number of < 50 colonies in a 2 ul spot. Each dilution was plated onto each of mMRS, supporting the growth of LAB and AAB, and mMRS supplemented with 10 mg/ml erythromycin and 10 mg/ml chloramphenicol, to inhibit the growth of LAB. The colony-forming units (CFUs) of both groups were manually or automatically counted (36) to compare the abundance of LAB and AAB in both the fly microbiome and diet. AAB and LAB are readily discriminated on MRS medium by their colony color: AAB are copper-colored, whereas the LAB are yellow or white (39). Fractional CFU abundances are shown as the ratio of the mean AAB:mean LAB abundance of all replicate measures with the ratio AAB:LAB in each sample shown as an individual point, and an overlayed violin plot to help visualize the distribution of the data. In some cases, the y-axis is shown on smaller than a 0 to 1 scale so that it is easier to see the difference in a minority population of LAB. Absolute CFU abundances are reported as the mean of all replicated measures, with standard error of the mean shown as error bars.

Differences in microbiota composition with variables in our experimental design were determined using PERMANOVA of a Bray-Curtis beta-diversity distance matrix. CFU counts were converted to an OTU table using custom R scripts and Bray-Curtis distances between samples were calculated using QIIME2 for both raw and rarefied CFU counts. CFU counts were rarefied to 2000 or 10000 CFUs, depending on the dataset. Differences in microbiota composition were defined using the two Bray-Curtis distance matrices and the Vegan R package (50). We interpreted significant variation in the raw and rarefied CFU counts as relating to variation in absolute and relative abundance in the microbiota, respectively. A Spearman correlation was used to test if there was a significant correlation between latitude and the mean LAB fraction of each geographic location. Significant differences between the absolute abundances of the LAB or AAB were determined using a Wilcoxon test or a Kruskal-Wallis test with a post-hoc Dunn test using the R packages dunn.test (51), rcompanion (52), and multcomp (53).

To discriminate between resident and transient microorganisms we analyzed siblings from the same vial at the time the flies were sorted and surface-sterilized on a CO_2_ pad. Some flies were directly homogenized and dilution-plated to measure the total internal microbiota. Some flies were briefly starved for 2 h in empty wells of a polystyrene 96-well plate, then transferred to a 96-well plate containing glass beads for homogenization and dilution plating. The starvation strategy eliminated transient microorganisms that could not persist for longer than the bulk flow of food, leaving behind resident microorganisms (e.g. (21, 34)). We estimated the abundance of the transient microorganisms in the flies as the difference between the average abundance of LAB and AAB in the unstarved and starved flies on a per-experiment basis (3 total experiments in time, each with 3 replicate vials of flies).

### *D. melanogaster* feeding preference for the microbiota

Microbial dietary preferences were determined in two conventionally-reared laboratory isofemale lines of *D. melanogaster* reared on a Y-G diet and using the FlyPad assay. The two fly lines were reported previously and were named F37 and ME89 and are called Fl and Me3 in this study (10). Adult female flies were briefly starved for 3 hours in clean, empty fly vials, and then introduced to the FlyPad arenas (54) as in our previous work (55). Each arena features two diet troughs that were prepared to contain 6.5 microliters of Y-G diet, except the preservative was omitted from the Y-G diet. In each arena the tough on the right was inoculated with 1 ml of OD_600_=0.1-normalized *A. orientalis* DmW_045 and the trough on the left side was inoculated with 1 ml of OD_600_=0.1-normalized *L. suionicum DmW_*098. The FlyPad arenas were placed in a lighted, 25°C, 75% RH incubator for 30 minutes to collect feeding preference data, where every time the flies stood on a platform and fed from the trough was read as a change in the system capacitance, or a ‘sip’. The number of times sips from each trough were taken was used as response variables in a generalized linear mixed effects model with a binomial family as in our previous analyses, with fly genotype as the main effect and experimental replicate as a random effect, using the lme4 R package (56). Four separate experiments were performed, each with 46, 8, 20, and 45 replicate arenas. The preference index is presented as the mean and s.e.m. of the linear prediction for each sample by the model.

### *D. melanogaster* feeding preference and microbiota composition

To examine the influence of fly feeding preferences on the microbiota composition of adult *D. melanogaster* we established an assay that blocked or enabled fly choice of their microbiota depending on how the available strains were mixed in the fly diet. We introduced 8 male and 8 female flies into an arena constructed from a modified 6-oz square-bottom fly bottle (VWR 75813-140) (the ‘top’) with a 35 mm petri dish (Greiner 627102) containing bacteria-inoculated Y-G diet (the ‘bottom’) (Fig. 1). The top was modified by drilling a 0.75-inch diameter hole into the bottom of the bottle and filling the hole with a Droso-Plug (Flystuff 59-200). The bottom was prepared by pouring 2.5 ml of Y-G diet into each half of a 35-mm petri dish, with the halves separated by a plastic square fit firmly between the sides of the dish to divide it in two; after the agar solidified the plastic piece was removed. Then, *A. orientalis* DmW_045 and *L. suionicum DmW_*098 were prepared as OD_600_ = 5 or 0.1 stocks, respectively (see Text S1), and inoculated to the plates in one of two treatments, each corresponding to a separated-condition or a mixed-condition treatment. In the separated condition, one-half of the diet plate was inoculated with 25 ml of DmW_045, and the other half was inoculated with 25 ml of DmW_098. In the mixed-condition treatment, flies were not given a choice between microbes because both halves of the diet were inoculated with 25 ml of a 1:1 homogeneous mixture of the two bacterial strains. The diet was incubated at 30°C for 24 hours before it was placed into the arena. Thus, between both treatments the flies were exposed to the same starting concentrations of microbes but with differences in their spatial separation. The assay began when a diet plate was placed at the bottom of a fly bottle containing the 16 flies and lasted for 3 days, during which the diet for each arena was replaced every 24 hours (3 times total) with a fresh diet plate prepared 24 hours prior as described above. After 3 days, whole-body flies from each arena were anesthetized with CO_2,_ surface-sterilized by brushing with 70% ethanol, and their microbiota composition was measured as described above. From each arena, three pairs of male flies and three pairs of female flies were homogenized and dilution plated.

We also collected diet samples from each partition in the arena by stabbing the diet partition three times with a wooden stick and spinning the stick in the 96-well plate containing beads and PBS to resuspend the collected material. We collected triplicate diet samples per diet partition in each arena. Unlike when we analyzed the fly microbiota, the diet data were first aggregated; meaning that the data were averaged by the diet condition bottle (which usually had 3-4 fly replicates or 3 separate sets of 3 ‘stabs’ into the diet). The replications for diet had to be aggregated by diet condition bottle because there was no justification for comparing one replicate from one side of the diet condition plate with any one replicate from the other side; each was equivalent and therefore needed to be averaged before they could be compared.

### Analysis of *D. melanogaster* starvation resistance

After 3 days of bacteria-treated diet exposure in our dietary preference assay, the adult flies were sorted on light CO_2_ anesthesia, and males and females were transferred to separate 30 ml vials containing 5 ml of 1% agarose. Their lifespan was monitored by recording fly mortality every 4 hours until all flies in a vial were dead. Significant differences in fly survival between the mixed- and separated-bacteria condition treatments were determined using a log-rank test in R (57–59), and compact letter displays were generated using the multcomp R package (53).

### Statistical analysis and data processing

Statistical analyses were performed in R using custom R scripts (File S1) and additional packages that were not cited above (60–66). Raw data files and a HTML-knitted version of the R Markdown script are available at https://github.com/johnchaston/microbiota2024/

## Results

### Wild fly lines from the eastern USA do not appear to vary in their genetic selection on the microbiota

To better understand how different wild fly populations select on their microbiota composition we compared microbiota composition between eight fly populations derived from collections in eight locations in the eastern USA and reared under gnotobiotic conditions (Fig. 2A). The relative abundances of AAB and LAB in the different populations were not significantly different from each other (F_7, 216_ = 2.06, R^2^ = 0.05, p = 0.054, Fig. 2B, Table S1). This finding conflicted with our previous observations, where the relative abundances of AAB and LAB in wild flies across the same latitudinal gradient significantly varied between locations (10). We reasoned that the wild conditions may have contributed to the differences in the microbiota composition of the wild flies. Therefore, we tested if the microbiota composition of these fly populations differed under conditions different between the lab and the wild: when flies could choose the microbes in their diets, when resident and transient microorganisms were considered separately, and when environmental humidity was elevated.

**Figure 2:**
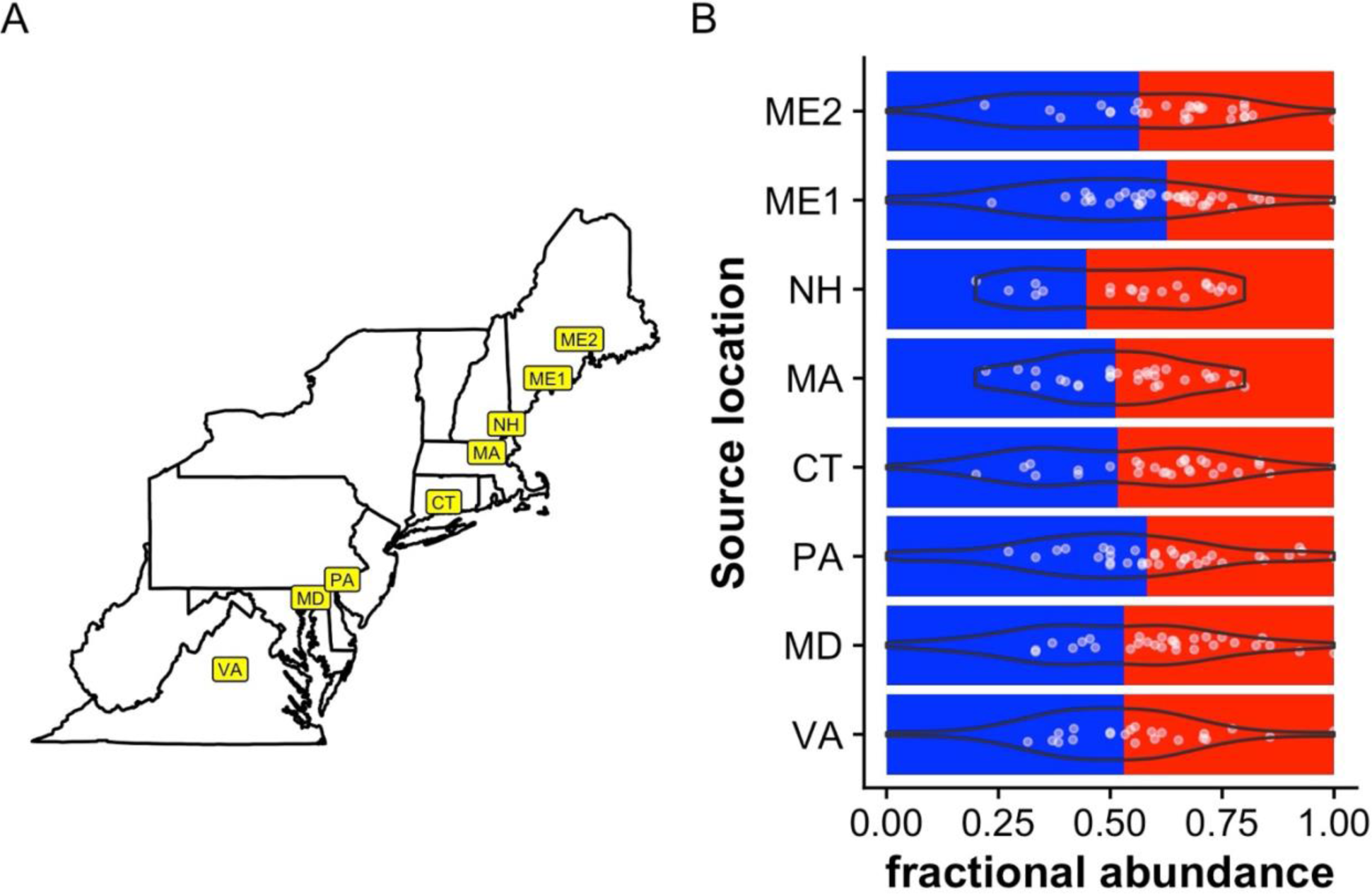
The relationship between fly collection site and microbiota composition. Microbiota composition was measured in gnotobiotic 6-sp common garden fly populations of isofemales lines originally collected from A) eight locations in the eastern USA. B) Fractional abundance of AAB (red) and LAB (blue), overlaid with violin plots and white points showing the fraction of LAB in each individual. Data are from 3 experiments, each targeting triplicate vials of flies per location (in some cases, the target number of vials was not reached because of low egg density). For each vial, 2 individual male and 2 individual female flies were homogenized.

### Fly choice of the microbiota promotes LAB abundance in the flies but is not geographically patterned

We reasoned that fly feeding preferences might have a greater impact on microbiota variation in the wild than the laboratory because while laboratory diets are well-mixed by larval activities, wild flies are likely to encounter diets with distinct microbial communities. Also, flies are known to have distinct feeding preferences for the microorganisms in their diets (40). We performed experiments under our own laboratory conditions that confirmed wild flies in the eastern USA have genotype-specific feeding preferences for microorganisms in their diets. Two wild fly genotypes derived from collections in Maine, USA (ME3) and Florida, USA (FL) each preferred diets inoculated with DmW_098 (Fig. 3A, Z_1, 119_ = -13.56, p < 10^-15^) when given a choice between *A. orientalis* DmW_045 and *L. suionicum* DmW_098. Thus, the flies have distinct feeding preferences. To test if fly feeding preferences can influence their microbiota composition, we exposed the flies to diets where the bacteria were mixed together or were spatially separated on the agar surface (Fig. 1). Conventionally reared flies bore a higher relative abundance of LAB when they fed on diets with separated, relative to mixed, bacterial conditions (F_1, 373_ = 10.93, R^2^ = 0.03, p < 0.001; Fig. 3B, Table S2). The diets where the bacteria were separated also bore significantly a significantly higher fraction of LAB than when the bacteria were mixed together (F_1, 77_ = 4.77, R^2^ = 0.06, p = 0.035, Fig. 3C, Table S3), but the fraction of LAB was 78-fold lower in the diets than the flies, demonstrating that the fly microbiota was not merely a reflection of the bacteria-conditioned diets (F_1, 231_ = 3.99, R^2^ = 0.02, p = 0.038, Fig. 3B-C, Table S4). Together, these findings demonstrate that conditions that permit flies to choose between the microbes in their diets alter the flies’ microbiota composition. However, the flies from different locations did not have significantly different microbiota composition in either the mixed (F_7, 191_ =1.74, R^2^ = 0.06, p = 0.08, Fig 3D, Table S5) or separated (F_7, 181_ = 0.59, R^2^ = 0.02, p = 0.80, Fig 3E, Table S6) condition arenas, showing that choosing the microbiota alone is not sufficient to lead to detectable, latitudinal variation in the fly microbiota composition.

**Figure 3.**
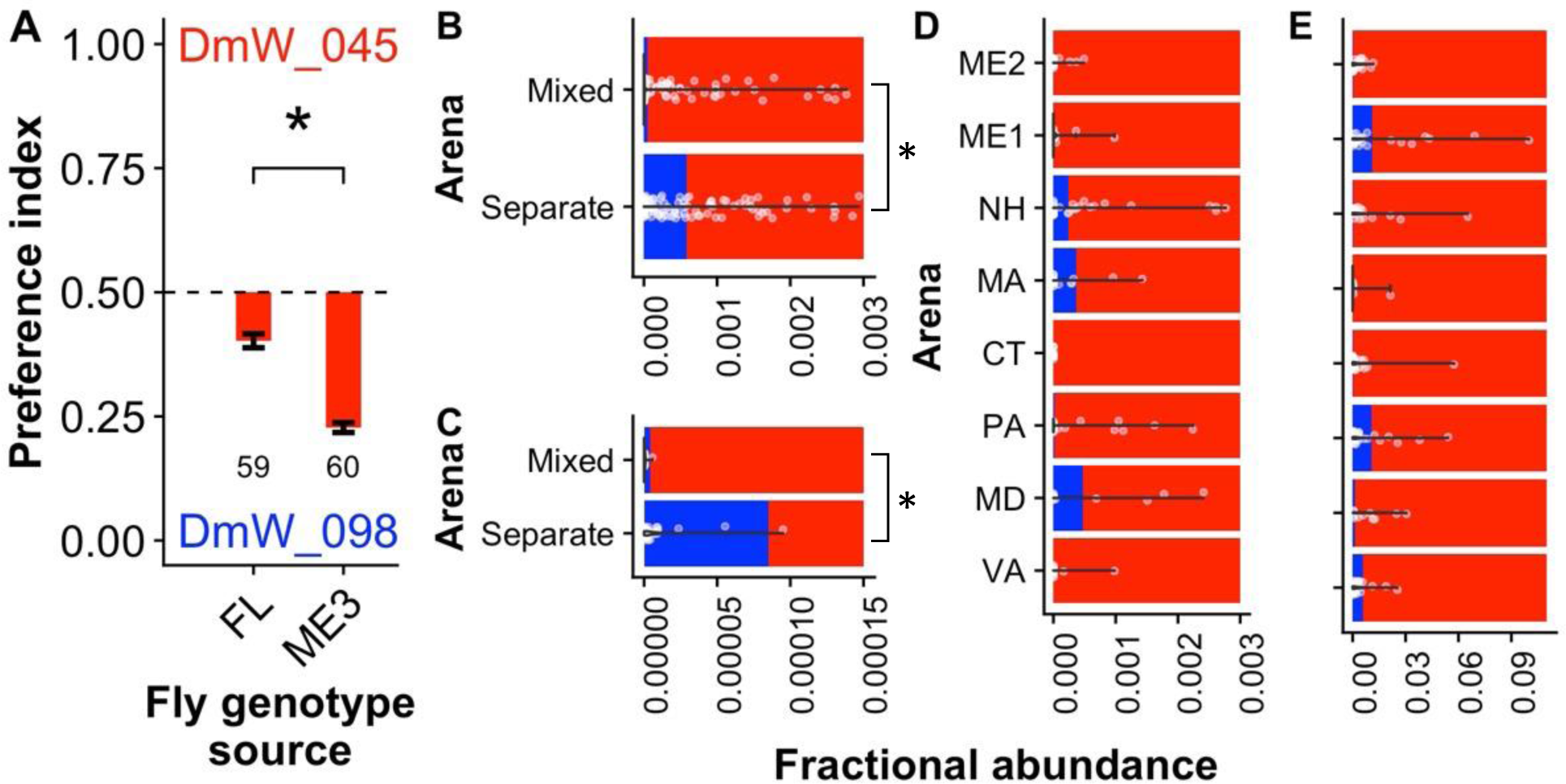
Bacteria-conditioned diets influence fly microbiota composition. A) Wild fly genotypes from the eastern USA (Florida (FL) and Maine (ME)), reared on Y-G diet and given a choice between *A. orientalis* DmW_045 and *L. suionicum DmW_*098. Numbers indicate the number of flies in the analysis, from across four separate experiments. The relative abundance of LAB and AAB in B) flies or C) diets after exposure to mixed- or separated-bacteria condition diets when all eight common garden populations were aggregated together. The relative abundance of LAB and AAB in flies from different population when reared in D) separated- or E) mixed-bacteria condition arenas. Different x-axis scales are used to facilitate seeing the LAB abundances, which were much lower in the mixed Bar color shows the bacterial type measured: AAB (red), LAB (Blue). Bars are overlaid with violin plots and white points showing the fraction of LAB in each individual. Data are from 2 experiments, each targeting triplicate vials of flies per location (in some cases, the target number of vials was not reached because of low egg density). From each vial, 3 pools of 2 male flies and 3 pools of 2 female flies were homogenized. * p < 0.05 by A) a generalized linear mixed effects model with a binomial family, or B-C) PERMANOVA.

### The resident and transient microbiota are distinct but do not vary in flies from different locations

To understand how different characteristics of the bacteria might influence natural variation in host colonization we tested if there is a relationship between the source location of the flies and the composition of the resident and transient portions of the fly microbiota. Residence of the microbiota is relevant to comparisons between the wild and the laboratory because flies are often unfed between collection and travel to the laboratory, during which time non-resident microorganisms are defecated. In the laboratory, we defined residents as microorganisms that colonized the flies after a 2 hour starvation period, and transients as the difference between microbial loads in starved flies and their unstarved, vial-matched siblings. There was a significant difference in the relative abundances of the resident and transient microbiota combined (unstarved flies) or resident bacteria only (starved flies), consistent with established work that a portion of the fly microbiota is transient (20, 21, 33, 34) (F_1, 389_ = 13.19, R^2^ = 0.03, p < 0.001, Fig. 4A, Table S7). However, the different fly lines did not have different abundances of the total (Fig. 1B, data collected in the same experiment as the rest of figure 3), or resident-only microbiota (F_7, 172_ = 1.44, R^2^ = 0.05, p =0.20, Fig. 4B, Table S8). Further, when we estimated the transient portion of the microbiota from the total and resident CFU counts, there was wide variation in the fractional abundance of LAB with source locations of the flies, but substantial overlap between data points, low overall N, and no visible trends with latitude (Fig. 4C). Taken together, we found no evidence that the fly genotypes selected for distinct relative abundances of their total, resident, or transient microbiota.

**Figure 4.**
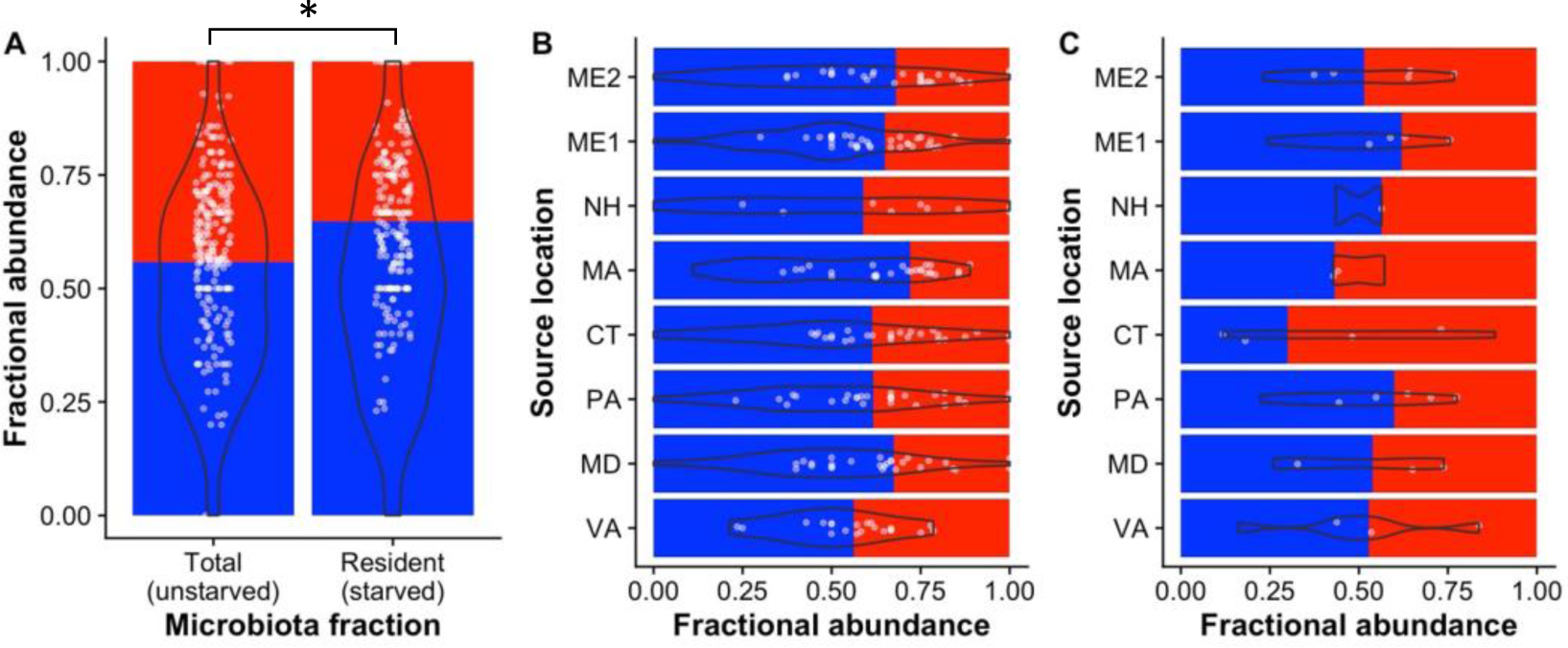
Differences in microbiota colonization of the flies. A) Relative abundance of total and resident bacteria in gnotobiotic 6-sp common garden fly populations of isofemales lines. Average differences in total bacteria between the population, collected in the same experiment are shown in Figure 1B. B) Average differences in the relative abudancen of resident microorganisms were measured in adult flies, starved for 2 hours. C) Transient microorganisms were calculated as the difference between the total and resident population. Data are shown as the fraction of experiment-wide means of AAB and LAB, overlaid with violin plots and white points showing the fraction of LAB in each individual. Data come from 3 experiments, each targeting triplicate vials of flies per location (in some cases, the target number of vials was not reached because of low egg density). For each vial, 2 individual male and 2 individual female flies were homogenized. * p < 0.05 by PERMANOVA.

### Humidity determines penetrance of a latitudinal pattern in the microbiota of flies from the eastern USA

Finally, we tested if the environment could influence fly genotype-dependent variation in microbiota composition. A major difference between the wild and laboratory environment was the humidity at which the laboratory experiments were conducted (25% RH in lab versus average daily RH between 70 and 83 % in our sampling locations; https-//nsrdb.nrel). Therefore, we tested if variation in humidity influenced the microbiota composition of the flies, including in the total and resident microbiota. Unlike at ambient humidity (Figs. 2B, 4B), fly populations reared at elevated humidities differed in the composition of their total (resident + transient; F_7, 453_ = 2.43, R^2^ = 0.03, p = 0.01, Table S9, Fig. 5A) and resident microbiota ( F_1, 379_ = 2.31, R^2^ = 0.03, p = 0.03, Table S10, Fig. 5B). Because the microbiota composition differed between the populations we tested if there was a correlation between the fraction of LAB in the flies and the latitude the flies were collected from. The relative abundance of the LAB was positively correlated with the source latitude of the flies in the total (S = 18, s = 0.79, p = 0.03) and transient (S =10, s = 0.88, p = 0.01, Fig. 5C), but not the resident (S = 106, s = -0.26, p = 0.54) microbiota. The significant positive correlation between relative abundance of LAB and the source latitude of the fly populations mirrors the pattern we previously observed in wild fly populations in the eastern USA (10) and provides evidence for a gradient in host genetic selection on the microbiota that was predicted by but not tested for in those previous analyses. These significant associations were observed when males and females were analyzed together, but were only apparent in the males when the sexes were analyzed separately (Table S11). Therefore, variation in humidity is sufficient to determine the penetrance of a sex-dependent gradient in host-genetic selection on the microbiota composition.

**Figure 5.**
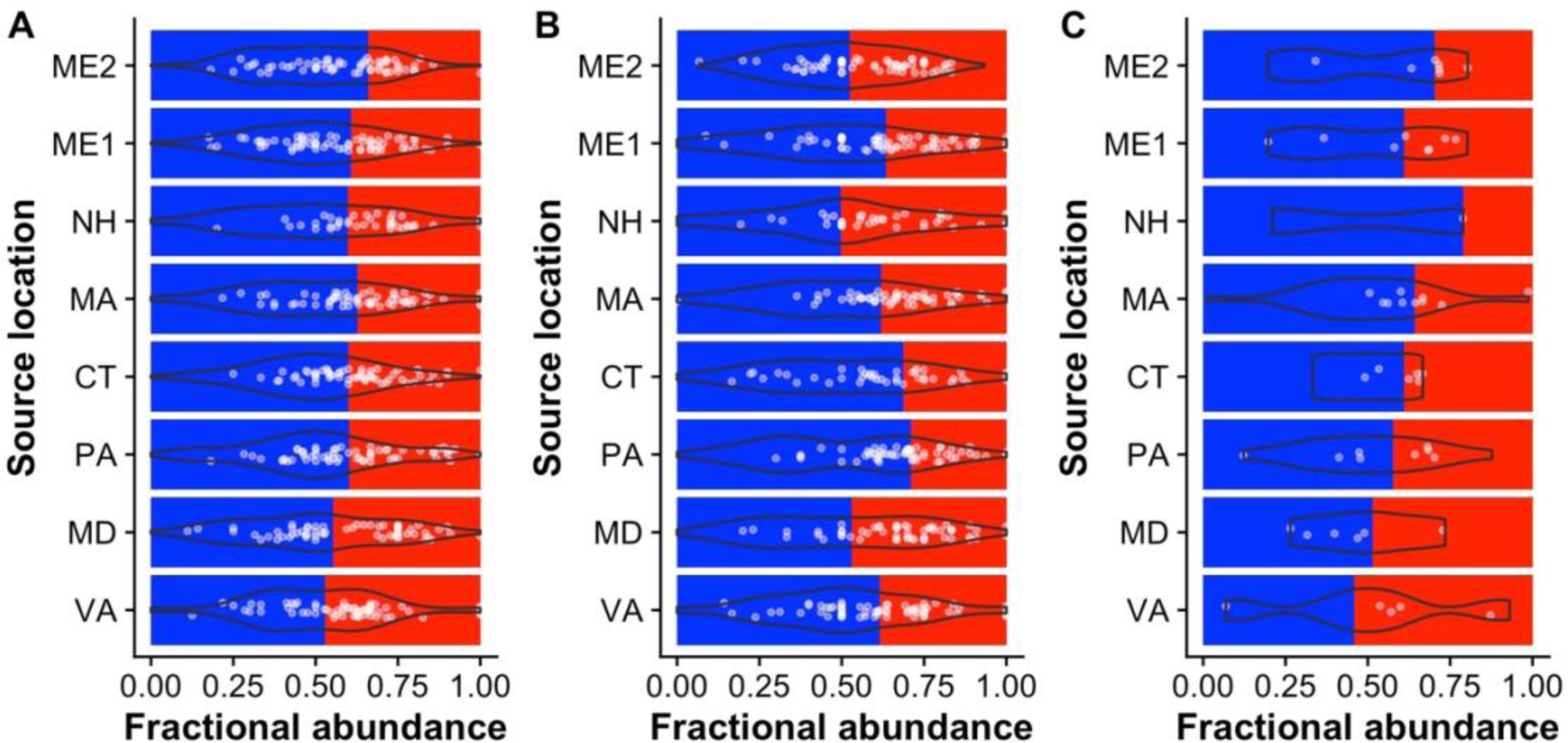
Humidity influences penetrance of a latitudinal gradient in host genetic selection on the microbiota. The A) total, B) resident, and C) transient microbiota composition of gnotobiotic 6-sp common garden fly populations of isofemales lines reared from the embryonic stage to 3 day old adulthood at 25% RH, then for 3 days at 50% and 60-75% humidity. Data are shown as the fraction of experiment-wide means of AAB and LAB, overlaid with violin plots and white points showing the fraction of LAB in each individual. Data come from 3 experiments, each targeting triplicate vials of flies per location (in some cases, the target number of vials was not reached because of low egg density). For each vial, 2 individual male and 2 individual female flies were homogenized.

### Fly choice-dependent variation in microbiota composition shapes a key fly life history trait

The experiments above document statistically significant but numerically small changes in microbiota composition under certain conditions. To contextualize the influence of relatively small changes in microbiota composition on the flies, we measured the starvation resistance of a laboratory strain of *D. melanogaster* after it was reared in the mixed- and separated-bacteria arenas for three days. Conventionally reared *D. melanogaster* in the separated-bacteria arenas survived longer than age-matched siblings in to the mixed-bacteria arenas (c^2^ _3, 579_ = 283, p < 10^-15^, Fig. 6, Table S12). These findings demonstrate that small (<1% relative abundance) changes in microbiota composition can be associated with significant changes in a biologically-relevant fly life history trait. These findings suggest that the relatively small changes in microbiota composition between different host genotypes, between flies that can choose their microbiomes, or in the different host niches occupied by the microbial populations are likely to have important influences on the adaptive traits of flies in the wild.

**Figure 6.**
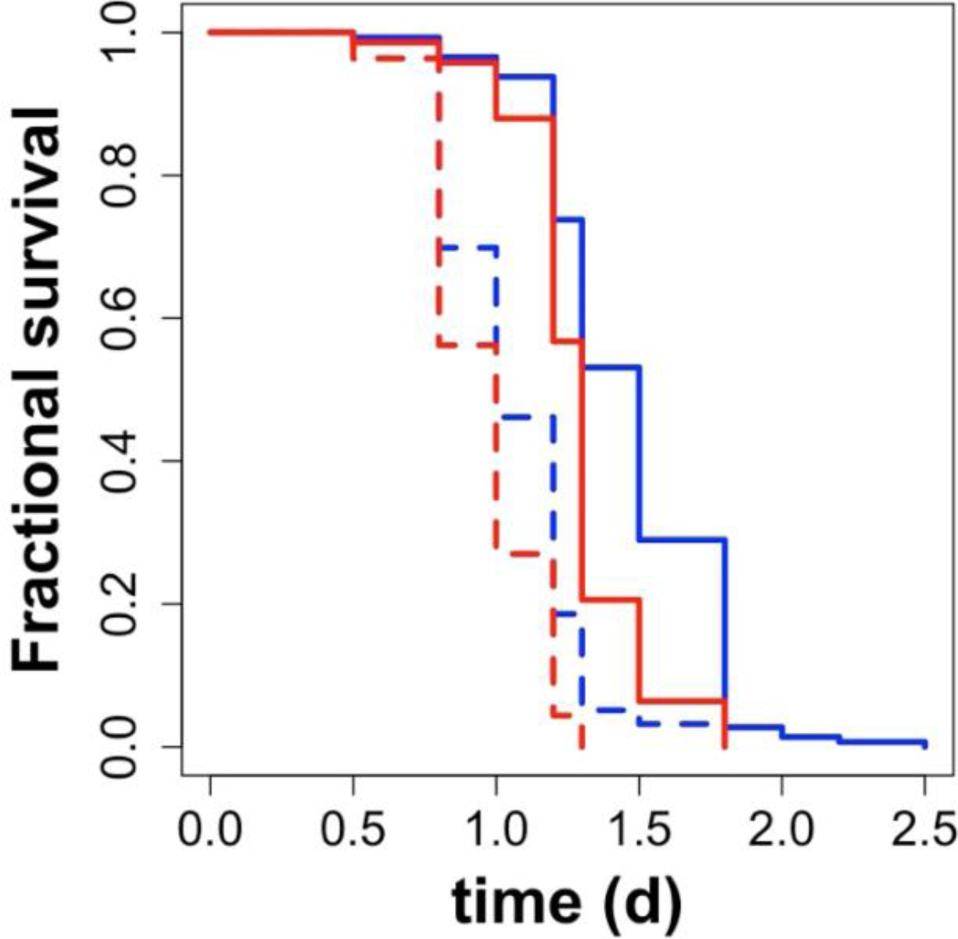
Small changes in microbiota composition are associated with significant variation in a key life history trait. Fractional survival of 4-6 days post eclosion male (dashed) and female (solid) flies from the mixed (red) or separated- (blue) bacteria condition when transferred to 1% agarose for a starvation resistance analysis.

## Discussion

Using wild flies collected from different locations in the USA we investigated the relationship between microbiota composition and host dietary preferences, the host niche where the bacteria localize, and humidity. Each of these factors influenced the microbiota composition of the flies, but we only detected differences in the microbiota composition of the different fly populations when humidity was elevated. Therefore, either the environmental condition of the flies is necessary for penetrance of the genetic variation in microbiota composition; or the colonization capacity of the bacteria and the acquisition preferences of the host are weaker than a low-RH suppression of host genetic variation in microbiota composition. Either explanation favors the idea that the environmental condition of humidity was the strongest determinant of microbiota composition among the factors we considered in this study. Taken together, the results emphasize there are key genotype-by-environment interactions that contribute to geographic patterns in host genetic selection on the microbiota.

Our analyses show that humidity contributes to the penetrance of the host genetic selection on microbiota composition. This is in addition to previous work that has already established diet and temperature as key environmental determinants of fly microbiota composition (19, 25, 27). Therefore, while these findings support the idea that genetic factors help shape microbial communities (9), they also reinforce that host genotype alone is insufficient to mediate specific patterns in microbiota composition in the wild. The insufficiency of host genotype alone to determine such patterns was previously shown by a sequencing analyses of the microbiota in flies from the eastern USA (32). In this study, the authors failed to detect a positive relationship between the fraction of LAB and the latitude from which flies were sampled, and detected low overall counts of LAB generally. Further, since these flies were sampled during humid periods, it seems unlikely that humidity alone is sufficient to lead wild flies to display the gradient in microbiota composition we observed (microbial sampling through the diet seems an obvious difference between our laboratory study and their wild sampling). These observations all support the current expectation that multiple factors contribute to patterns of microbiota composition in the wild.

In addition to focusing on the ratio between AAB and LAB in adult flies in the main text, the supporting text reports on the variation in absolute abundance of bacteria in flies (Text S2). We focused on relative abundance to draw conclusions between this work and our previous sequencing study, which made no conclusions about absolute abundance. However, absolute abundance of the microbiota in *D. melanogaster* should not be ignored as it can determine some of the microbiota influence in the flies. For example, absolute abundance of AAB is negatively correlated with the fat content of the flies, presumably because larger bacterial loads can consume more glucose from the fly diet and fewer nutritional resources are consequently available to the flies (9, 46). There is a known positive correlation between body size and latitude (67–70) in flies collected from the eastern USA, and we therefore had expected there might be a positive correlation between bacterial load and the source latitude of the flies reported in this study. However, across all of our treatments, we never detected a significant correlation between the source location of the flies and AAB abundance, LAB abundance, or total bacterial abundance in the flies. We did not measure the body size of the flies in this study, but if they vary in size as expected, our microbiota abundance data would suggest that variation in body size is not necessarily correlated with bacterial loads in the flies. Therefore, either gut size does not strictly correlate with body size; or factors beyond gut size help determine bacterial load in the flies. There is existing support for the latter idea, for example, the numerous indications that host immunity influences the fly microbiota composition (recently, (71–75)) and that most of the living bacteria occupy a distinct niche in the foregut, not the gut (36). Relative to the host niche of the bacteria, our finding that the relative abundance of the transient, but not the colonizing, microbiota varies with latitude suggests that latitudinal variation is driven by variation outside of the colonized foregut niche. The reciprocal corollary of these observations is that the foregut niche microbiota displays little variation with geography and may be conserved across a variety of host populations; if so, the strong conservation would suggest an as yet undescribed selective benefit of maintaining a particular ratio of microorganisms.

The bacteria-conditioned diet experiment provided an unexpected opportunity to test if different abundances of the same bacterial community could influence host phenotypes. Previous work has shown that inoculating the same host with different starting ratios of microorganisms (46), or rearing hosts at varying densities (76) can also significantly alter the fly microbiota composition of the same starting sets of microorganisms. In our experiments, the change in microbiota composition were less than a 1% shift in relative abundance of LAB across any of the starting microbial treatments. Despite the small magnitude of the change in microbiota composition, the survival of laboratory flies under starvation conditions shifted dramatically with the microbial treatment. If the absolute abundance of the AAB increased in the mixed-bacteria condition versus the separated-bacteria condition, we would have expected much poorer starvation survival in the mixed-bacteria condition: more AAB predicts lower glucose content in the diet, less fat storage in the flies, and a shorter period of starvation resistance (10, 15, 46, 77). Instead, the increased starvation survival of the flies in the separated bacteria condition appears to have been driven by the LAB because there was no change in the absolute abundance of AAB in the flies; however, we cannot rule out that a change in AAB gene expression or protein function differed between the two diet conditions; or it may be possible that the effect is related to dietary manipulation by the bacteria during the pre-condition stage (77). Regardless, the dramatic shift in host starvation survival shows that having the capacity to genetically select the microbiota even in small ways could have substantial life history impacts.

In conclusion, the intricate interplay between host genetics, environment, and microbiota composition remains a focal point of investigation in understanding the complex host-microbe interactions. We revealed a nuanced relationship between fly genotype and microbiota composition, with factors such as diet preferences, environment, and starting microbial composition contributing to the observed patterns. Our findings underscore the complexity of microbiota dynamics, highlighting how plastic changes in the adult fly environment (humidity) can override patterns established from birth. While host genetic influences are evident, their manifestation is context-dependent, necessitating consideration of various factors. This work advances our understanding of microbiota-host interactions and emphasizes the interplay between host genetic and environmental influences on animal-associated microbial communities.

## Acknowledgements

Research reported in this publication was supported in part by the National Institute ofGeneral Medical Sciences of the National Institutes of Health under Award Number R15GM140388. The content is solely the responsibility of the authors and does not necessarily represent the official views of the National Institutes of Health.

