## Supporting information for "Humidity determines penetrance of a latitudinal gradient in genetic selection on the microbiota by *Drosophila melanogaster*"

### Supporting tables

**Table S1. PERMANOVA results for CFU counts (AAB and LAB) of gnotobiotic flies in eight populations originally collected from each of eight different locations in the eastern USA.** See Figure 2B.

| | df [a] | SS [b] | $R^2$ | F | p [c] | |
|---|---|---|---|---|---|---|
| *Geography* | 7 | 0.33 | 0.05 | 2.06 | 0.054 | . |
| *Sex* | 1 | 0.06 | 0.01 | 2.70 | 0.11 | |
| *Exp* [d] | 2 | 0.10 | 0.02 | 2.24 | 0.11 | |
| *Plate* [e] | 7 | 0.33 | 0.05 | 2.05 | 0.04 | * |
| *G * S* [f] | 7 | 0.06 | 0.01 | 0.36 | 0.92 | |
| *Plate / Vial* [g] | 90 | 3.33 | 0.51 | 1.63 | 0.01 | * |
| *Residual* | 102 | 2.31 | 0.35 | | | |
| *Total* | 216 | 6.52 | 1.00 | | | |

[a] degrees of freedom
[b] sum of squares
[c] p-value
[d] One of three separate experiments in time
[e] 96-well plate on which individual replicates of flies were homogenized and dilution plated;
[f] "*" = the interaction term
[g] the source vial the flies were grown in (30-50 mixed sex flies per vial); "/" = the nesting term

**Table S2. PERMANOVA results for CFU counts (AAB and LAB) of gnotobiotic flies from the eight populations in our study, exposed to mixed- and separated-bacteria diet conditions.** See Figure 3B.

| | *df* [a] | *SS* [b] | $R^2$ | *F* | *p* [c] | |
|---|---|---|---|---|---|---|
| *Arena* [d] | 1 | 0.01 | 0.03 | 10.93 | 0.001 | *** |
| *Sex* | 1 | 0.00 | 0.01 | 2.94 | 0.081 | . |
| *Geography* | 7 | 0.00 | 0.01 | 0.57 | 0.805 | |
| *Exp* [e] | 1 | 0.00 | 0.01 | 2.52 | 0.114 | |
| *A * S* [f] | 1 | 0.00 | 0.01 | 2.99 | 0.091 | . |
| *E / Plate* [g] | 2 | 0.00 | 0.01 | 1.87 | 0.160 | |
| E / P / Arena [h] | 69 | 0.06 | 0.16 | 0.87 | 0.606 | |
| Residual | 291 | 0.27 | 0.77 | | | |
| *Total* | 373 | 0.36 | 1.00 | | | |

[a] degrees of freedom
[b] sum of squares
[c] p-value
[d] mixed- or separated-bacteria condition
[e] One of two separate experiments in time
[f] * = the interaction term
[g] 96-well plate on which individual replicates of flies were homogenized and dilution plated; / represents the nesting term
[h] the source arena the flies were collected from

**Table S3. PERMANOVA results for CFU counts (AAB and LAB) of the third-day mixed- and separated-bacteria diet conditions, after 24 hours of exposure to flies.** The data were aggregated by diet condition bottle, unlike in Table S2 where raw counts for flies were used (see methods for explanation why aggregation was necessary). See Figure 3C.

| | ***df*** [a] | ***SS*** [b] | ***$R^2$*** | ***F*** | ***p*** [c] | |
|---|---|---|---|---|---|---|
| *Arena* [d] | 1 | 0.00 | 0.06 | 4.77 | 0.03 | * |
| *Geography* | 7 | 0.00 | 0.10 | 1.18 | 0.31 | |
| *Exp* [e] | 1 | 0.00 | 0.03 | 2.38 | 0.14 | |
| *A * G* | 7 | 0.00 | 0.06 | 0.65 | 0.68 | |
| Exp / Plate [g] | 2 | 0.00 | 0.03 | 1.33 | 0.28 | |
| Residual | 59 | 0.00 | 0.72 | | | |
| *Total* | 77 | 0.00 | 1.00 | | | |

[a] degrees of freedom
[b] sum of squares
[c] p-value
[d] mixed- or separated-bacteria condition
[e] One of two separate experiments in time
[f] * = the interaction term
[g] 96-well plate on which individual replicates of flies were homogenized and dilution plated; / represents the nesting term

**Table S4. PERMANOVA results for CFU counts (AAB and LAB) of flies and the third-day mixed- and separated-bacteria diet conditions.** A single analysis to compare the microbiota composition between panels 3B and 3C.

| | *df* [a] | *SS* [b] | $R^2$ | *F* | *p* [c] | |
|---|---|---|---|---|---|---|
| *Arena* [d] | 1 | 0.0018 | 0.03 | 7.25 | 0.001 | *** |
| *Geography* | 7 | 0.0016 | 0.03 | 0.91 | 0.469 | |
| *Type* [e] | 1 | 0.0010 | 0.02 | 3.99 | 0.038 | * |
| *Exp* [f] | 1 | 0.0003 | 0.01 | 1.35 | 0.277 | |
| *A * G* [g] | 7 | 0.0017 | 0.03 | 0.95 | 0.482 | |
| *A * T* | 1 | 0.0009 | 0.02 | 3.74 | 0.043 | * |
| G * T | 7 | 0.0009 | 0.01 | 0.50 | 0.758 | |
| E / Plate [h] | 5 | 0.0008 | 0.01 | 0.62 | 0.651 | |
| A * G * T | 7 | 0.0008 | 0.01 | 0.48 | 0.783 | |
| Residual | 194 | 0.0490 | 0.83 | | | |
| *Total* | 231 | 0.0589 | 1.00 | | | |

[a] degrees of freedom
[b] sum of squares
[c] p-value
[d] mixed- or separated-bacteria condition
[e] flies or diets
[f] One of two separate experiments in time
[g] * = the interaction term
[h] plate = 96-well plate on which individual replicates of flies were homogenized and dilution plated; / represents the nesting term

**Table S5. PERMANOVA results for CFU counts (AAB and LAB) of flies in the mixed bacteria arenas.** See Figure 3D.

| | ***df*** [a] | ***SS*** [b] | $R^2$ | ***F*** | ***p*** [c] | |
|---|---|---|---|---|---|---|
| *Geography* | 7 | 0.00 | 0.06 | 1.74 | 0.08 | . |
| *Sex* | 1 | 0.00 | 0.00 | 0.01 | 0.93 | |
| *Exp* [d] | 1 | 0.00 | 0.00 | 0.75 | 0.44 | |
| *G * S* [e] | 7 | 0.00 | 0.01 | 0.39 | 0.92 | |
| *E / Plate* [f] | 3 | 0.00 | 0.01 | 0.89 | 0.46 | |
| *E / P / Arena* [g] | 28 | 0.00 | 0.22 | 1.67 | 0.07 | . |
| *Residual* | 144 | 0.00 | 0.69 | | | |
| *Total* | 191 | 0.00 | 1.00 | | | |

[a] degrees of freedom
[b] sum of squares
[c] p-value
[d] One of two separate experiments in time
[e] * = the interaction term
[f] plate = 96-well plate on which individual replicates of flies were homogenized and dilution plated; / represents the nesting term
[f] Arena = the source arena flies were collected from, one of three arenas per condition in each experiment

**Table S6. PERMANOVA results for CFU counts (AAB and LAB) of flies in the separated bacteria arenas.** See Figure 3F.

| | *df* [a] | *SS* [b] | $R^2$ | *F* | *p* [c] | |
|---|---|---|---|---|---|---|
| *Geography* | 7 | 0.01 | 0.02 | 0.59 | 0.80 | |
| *Sex* | 1 | 0.01 | 0.02 | 2.90 | 0.10 | . |
| *Exp* [d] | 1 | 0.00 | 0.01 | 1.92 | 0.18 | |
| *G * S* [e] | 7 | 0.01 | 0.02 | 0.51 | 0.82 | |
| *E / Plate* [f] | 3 | 0.01 | 0.02 | 1.20 | 0.30 | |
| *E / P / Arena* [g] | 29 | 0.05 | 0.14 | 0.87 | 0.56 | |
| *Residual* | 133 | 0.27 | 0.76 | | | |
| *Total* | 181 | 0.36 | 1.00 | | | |

[a] degrees of freedom
[b] sum of squares
[c] p-value
[d] One of two separate experiments in time
[e] * = the interaction term
[f] plate = 96-well plate on which individual replicates of flies were homogenized and dilution plated; / represents the nesting term
[g] Arena = the source arena flies were collected from, one of three arenas per condition in each experiment

**Table S7. PERMANOVA results for CFU counts (AAB and LAB) of the total and resident-only portions of gnotobiotic flies from the eight populations in our study.** See Figure 4A.

| | *df* [a] | *SS* [b] | $R^2$ | *F* | *p* [c] | |
|---|---|---|---|---|---|---|
| Geography | 7 | 0.34 | 0.03 | 2.21 | 0.036 | * |
| Sex | 1 | 0.02 | 0.00 | 1.04 | 0.306 | |
| Niche [d] | 1 | 0.29 | 0.03 | 13.19 | 0.001 | *** |
| Exp [e] | 2 | 0.22 | 0.02 | 4.91 | 0.008 | ** |
| Plate [f] | 15 | 0.46 | 0.04 | 1.39 | 0.164 | |
| G * S [g] | 7 | 0.06 | 0.01 | 0.38 | 0.926 | |
| G * N | 7 | 0.42 | 0.04 | 2.66 | 0.007 | ** |
| S * N | 1 | 0.04 | 0.00 | 1.65 | 0.206 | |
| Plate / Vial [h] | 175 | 6.04 | 0.51 | 1.55 | 0.003 | ** |
| Residual | 173 | 3.86 | 0.33 | | | |
| *Total* | 389 | 11.76 | 1.00 | | | |

[a] degrees of freedom
[b] sum of squares
[c] p-value
[d] flies in the directly-homogenized (total microbes) or briefly-starved (only resident microbes) treatments
[e] One of three separate experiments in time
[f] 96-well plate on which individual replicates of flies were homogenized and dilution plated
[g] "*" = the interaction term
[h] "/" = the nesting term; the source vial the flies were grown in (30-50 mixed sex flies per vial)

**Table S8. PERMANOVA results for CFU counts (AAB and LAB) of the resident-only portion of gnotobiotic flies from the eight populations in our study.** See Figure 4B.

| | *df* [a] | *SS* [b] | $R^2$ | *F* | *p* [c] | |
|---|---|---|---|---|---|---|
| Geography | 7 | 0.23 | 0.05 | 1.44 | 0.19 | |
| Sex | 1 | 0.00 | 0.00 | 0.04 | 0.82 | |
| Exp [d] | 2 | 0.18 | 0.04 | 3.90 | 0.02 | * |
| Plate[e] | 6 | 0.27 | 0.05 | 1.97 | 0.07 | . |
| G * S [f] | 7 | 0.13 | 0.03 | 0.83 | 0.56 | |
| Plate / Vial [g] | 78 | 2.55 | 0.52 | 1.45 | 0.05 | * |
| Residual | 71 | 1.60 | 0.32 | | | |
| *Total* | 172 | 4.95 | 1.00 | | | |

[a] degrees of freedom
[b] sum of squares
[c] p-value
[d] One of three separate experiments in time
[e] 96-well plate on which individual replicates of flies were homogenized and dilution plated
[f] "*" = the interaction term
[g] "/" = the nesting term; the source vial the flies were grown in (30-50 mixed sex flies per vial)

**Table S9. PERMANOVA results for CFU counts (AAB and LAB) of the total microbiota in gnotobiotic flies from the eight populations in our study when reared at 50% and 60-75% RH.** See Figure 5A.

| | *df* [a] | *SS* [b] | $R^2$ | *F* | *p* [c] | |
|---|---|---|---|---|---|---|
| *Geography* | 7 | 0.42 | 0.03 | 2.43 | 0.014 | * |
| *Sex* | 1 | 0.24 | 0.02 | 9.81 | 0.004 | ** |
| *Humidity* [d] | 1 | 0.08 | 0.01 | 3.23 | 0.083 | . |
| *Exp* [e] | 2 | 0.29 | 0.02 | 5.91 | 0.003 | ** |
| *Plate* [f] | 9 | 0.44 | 0.03 | 1.97 | 0.051 | . |
| *G * S* [g] | 7 | 0.16 | 0.01 | 0.92 | 0.501 | |
| *G * H* | 7 | 0.34 | 0.02 | 1.97 | 0.058 | . |
| *S * H* | 1 | 0.02 | 0.00 | 1.00 | 0.320 | |
| *Plate / Vial* [h] | 186 | 6.45 | 0.45 | 1.39 | 0.011 | * |
| *Residual* | 232 | 5.78 | 0.41 | | | |
| *Total* | 453 | 14.25 | 1.00 | | | |

[a] degrees of freedom
[b] sum of squares
[c] p-value
[d] flies reared at ambient (~25%) RH, at 50% RH, or at 60-75% RH
[e] One of three separate experiments in time
[f] 96-well plate on which individual replicates of flies were homogenized and dilution plated
[g] "*" = the interaction term
[h] "/" = the nesting term; vial = the source vial the flies were grown in (30-50 mixed sex flies per vial)

**Table S10. PERMANOVA results for CFU counts (AAB and LAB) of the resident microbiota in gnotobiotic flies from the eight populations in our study when reared at 50% and 60-75% RH.** See Figure 5B.

| | ***df*** [a] | ***SS*** [b] | $R^2$ | ***F*** | ***p*** [c] | |
|---|---|---|---|---|---|---|
| *Geography* | 7 | 0.45 | 0.03 | 2.31 | 0.028 | * |
| *Sex* | 1 | 0.24 | 0.02 | 8.55 | 0.009 | ** |
| *Humidity* | 1 | 0.12 | 0.01 | 4.45 | 0.046 | * |
| *Exp* | 2 | 0.42 | 0.03 | 7.50 | 0.001 | *** |
| *Plate* | 8 | 0.37 | 0.03 | 1.64 | 0.127 | |
| *G * S* | 7 | 0.31 | 0.02 | 1.61 | 0.137 | |
| *G * H* | 7 | 0.25 | 0.02 | 1.26 | 0.287 | |
| *S * H* | 1 | 0.01 | 0.00 | 0.49 | 0.478 | |
| *Plate / Vial* | 170 | 6.13 | 0.47 | 1.29 | 0.055 | |
| *Residual* | 175 | 4.88 | 0.37 | | | |
| *Total* | 379 | 13.18 | 1.00 | | | |

[a] degrees of freedom
[b] sum of squares
[c] p-value
[d] flies reared at ambient (~25%) RH, at 50% RH, or at 60-75% RH
[e] One of three separate experiments in time
[f] 96-well plate on which individual replicates of flies were homogenized and dilution plated
[g] "*" = the interaction term
[h] "/" = the nesting term; vial = the source vial the flies were grown in (30-50 mixed sex flies per vial)

**Table S11. The correlation between latitude and absolute or relative abundance of the microbiota in the 8 fly populations used in this study, when reared gnotobiotic at elevated humidity**.

Total

| | *Male & Female* | | | *Male-only* | | | *Female-only* | | |
|---|---|---|---|---|---|---|---|---|---|
| | $S^a$ | $\sigma^b$ | $p^c$ | $S$ | $\sigma$ | $p$ | $S$ | $\sigma$ | $p$ |
| *Total (absolute)* | 46 | 0.45 | 0.27 | **20** | **0.76** | **0.04** | 84 | 0 | 1 |
| *AAB (absolute)* | 84 | 0 | 1 | 44 | 0.48 | 0.24 | 92 | -0.1 | 0.84 |
| *LAB (absolute)* | 32 | 0.62 | 0.11 | **18** | **0.79** | **0.03** | 76 | 0.1 | 0.84 |
| *Relative* | **18** | **0.79** | **0.03** | **12** | **0.86** | **0.01** | 44 | 0.48 | 0.24 |

[a] S statistic
[b] Spearman's rho;
[c] p-value.

Resident

| | $S^a$ | $\sigma^b$ | $p^c$ | $S$ | $\sigma$ | $p$ | $S$ | $\sigma$ | $p$ |
|---|---|---|---|---|---|---|---|---|---|
| *Total (absolute)* | 84 | 0 | 1 | 88 | -0.05 | 0.93 | 96 | -0.14 | 0.75 |
| *AAB (absolute)* | 88 | -0.05 | 0.93 | 88 | -0.05 | 0.93 | 76 | 0.1 | 0.84 |
| *LAB (absolute)* | 88 | -0.05 | 0.93 | 118 | -0.4 | 0.33 | 96 | -0.14 | 0.75 |
| *Relative* | 106 | -0.26 | 0.54 | 110 | -0.31 | 0.46 | 94 | -0.12 | 0.79 |

[a] S statistic
[b] Spearman's rho;
[c] p-value.

Transient

| | *Male & Female* | | | *Male-only* | | | *Female-only* | | |
|---|---|---|---|---|---|---|---|---|---|
| | $S^a$ | $\sigma^b$ | $p^c$ | $S$ | $\sigma$ | $p$ | $S$ | $\sigma$ | $p$ |
| *Total (absolute)* | 42 | 0.50 | 0.22 | 48 | 0.43 | 0.30 | 52 | 0.07 | 0.91 |
| *AAB (absolute)* | 72 | 0.14 | 0.75 | 54 | 0.36 | 0.39 | 56 | 0.00 | 1.00 |
| *LAB (absolute)* | 22 | 0.74 | 0.05 | 28 | 0.67 | 0.08 | 32 | 0.43 | 0.35 |
| *Relative* | **10** | **0.88** | **0.01** | **14** | **0.83** | **0.02** | 14 | 0.75 | 0.07 |

[a] S statistic
[b] Spearman's rho;
[c] p-value.

**Table S12. Log-rank test statistics for Figure 6 data**.

| | *N* | *Observed (O)* | *Expected (E)* | *(O-E)^2/E* | *(O-E)^2/V* |
|---|---|---|---|---|---|
| *Female Separated* | 145 | 145 | 246.2 | 41.6 | 135.46 |
| *Male Separated* | 156 | 156 | 105.2 | 24.6 | 46.95 |
| *Female Mixed* | 141 | 141 | 167.2 | 4.1 | 9.54 |
| *Male Mixed* | 137 | 137 | 60.4 | 97 | 157.84 |

**Text S1**

**Optimizing the bacteria-conditioned diet experiments**

To test if fly preferences for microbes could alter their microbiota composition, we set up a 72-hour test condition that would permit or restrict fly choice for dietary microbiota by administering the microorganisms in shared or separate dietary compartments in a closed arena (Fig. 1). In addition to measuring microbiota composition in the flies placed in the arenas, we also quantified the visual cue of egg laying on the plates and performed CFU plating of diets. The prospective analysis of egg laying provided clues that fly movement or other conditions in the arena did not lead to rapid mixing of the microbiota between partitions of the mixed-bacteria condition arena. In our first efforts, we inoculated the diets in the arenas with the six bacterial species listed in the main text, each normalized to $OD_{600} = 0.1$, mixed in equal ratios, and inoculated to the diet in 25 μl aliquots. For the mixed-bacteria condition, all six strains were mixed together and 25 μl was inoculated to each of two sides of the plate. For the separated-bacteria condition, the three LAB strains were mixed together and inoculated to one random half of the plate, and the three AAB strains were mixed together and inoculated to the other half of the plate, each in a 25 μl volume. For the separated-bacteria condition plate, it did not matter which half of the plate had LAB or AAB because the plates could freely rotate in the arena. Consistent with known *D. melanogaster* preference to lay eggs on AAB-inoculated surfaces (1), *D. melanogaster* flies collected from three locations in the eastern United States (Florida and Maine, same locations as the Fl and Me3 locations reference in the main text, as well as a 3rd location in Media, Pennsylvania) laid significantly fewer eggs on the LAB-inoculated side of the separated-bacteria condition diet plates after 24 hours than on the AAB-inoculated side of these plates (Fig. S1AC, $W_{2, 125} = 3182$, $p < 10^{-8}$). Conversely, there was no significant difference between either the number of eggs laid on either side of the mixed-bacteria condition plates (Fig S1BD, $W_{2, 126} = 1938$, $p = 0.82$), or the total number of eggs laid on diet plates inoculated under the choice or control condition (Fig S1E, $W_{2, 125} = 2239$, $p = 0.22$). The egg laying patterns confirm that the inoculation strategy for the mixed- or separated-bacteria conditions did not alter total fly reproductive output, a major indicator of fly health and productivity, but did influence fly oviposition, a key fly behavior within the arena. We proceeded to analyze the microbiota composition of wild fly lines from the eight locations in the eastern USA, which displayed similar qualitative patterns in egg laying as these three wild populations.

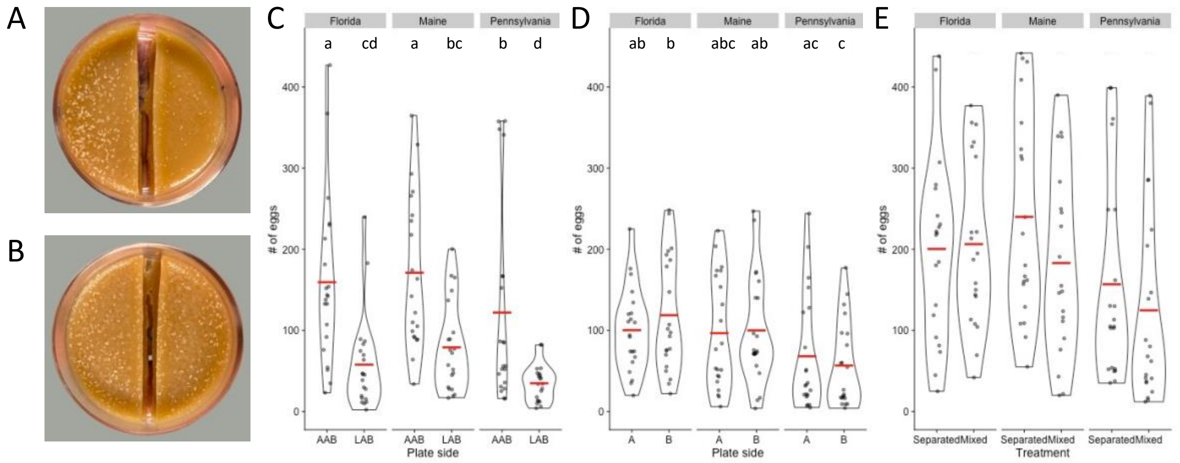


**Figure S1**. Representative images showing eggs laid by CantonS flies on A) separated- or B) mixed-condition bacteria plates. We counted eggs from the plates used in three sequential days in a single experiment for C) the AAB- or LAB- inoculated sides of separated bacteria condition diet plates; D) randomly assigned "A" or "B" inoculated sides of mixed bacteria condition diet plates; E) both sides of either separated- or mixed-bacteria condition plates. Three different isofemale lines were tested for each condition, each originally collected from a different location in the eastern USA. Each point represents a count from a single plate. Letters over the bars report statistically significant differences in the counts for each genotype determined by a Kruskal-Wallis test, with a post-hoc Dunn test if the differences were significant.
A: $\chi^2_{5, 126} = 46.91$, p = 0
B: $\chi^2_{5, 126} = 15.97$, p = 0.01
C: $\chi^2_{5, 126} = 1,54$, p = 0.21

Analysis of the fly microbiota of the eight wild collected populations after three days in the mixed- and separated- condition arenas revealed a low overall count of LAB (< 0.5% overall), and a significant effect of fly sex (Fig. S2A; $F_{1, 343} = 4.45$, $R^2 = 0.01$, $p = 0.04$) but not the bacteria-conditioned diet arenas (Fig. S2B, $F_{1, 343} = 3.26$, $R^2 = 0.01$, $p = 0.08$) or fly geography ($F_{7, 343} = 1.62$, $R^2 = 0.03$, $p = 0.14$) on microbiota composition (Table S13). Thus, giving the flies the opportunity to choose their microbiota did not significantly influence the ratio of LAB to AAB. There was significant variation in the absolute CFU counts with each of fly sex (Fig. S2C, $F_{1, 463} = 27.42$, $R^2 = 0.05$, $p < 0.01$), arena condition (Fig. S2D, $F_{1, 463} = 4.986$, $R^2 = 0.01$, $p = 0.004$), and fly source location (Fig. S2E, $F_{7, 463} = 3.51$, $R^2 = 0.04$, $p < 0.001$) (Table S13). Also, very few LAB overall were detected. The low overall counts of LAB and lack of a significant effect of the treatment raised questions about how we might increase the relative abundance of LAB in the flies and if doing so would lead to a significant effect of fly choice on microbiota composition. The data also pointed to the need to determine if the microbiota composition of two diet conditions varied as a control for understanding variation in fly microbiota composition. We began these efforts by measuring microbiota composition in the flies and diets when the inoculum density of LAB to the bacteria-conditioned diet arenas was increased 50-fold.

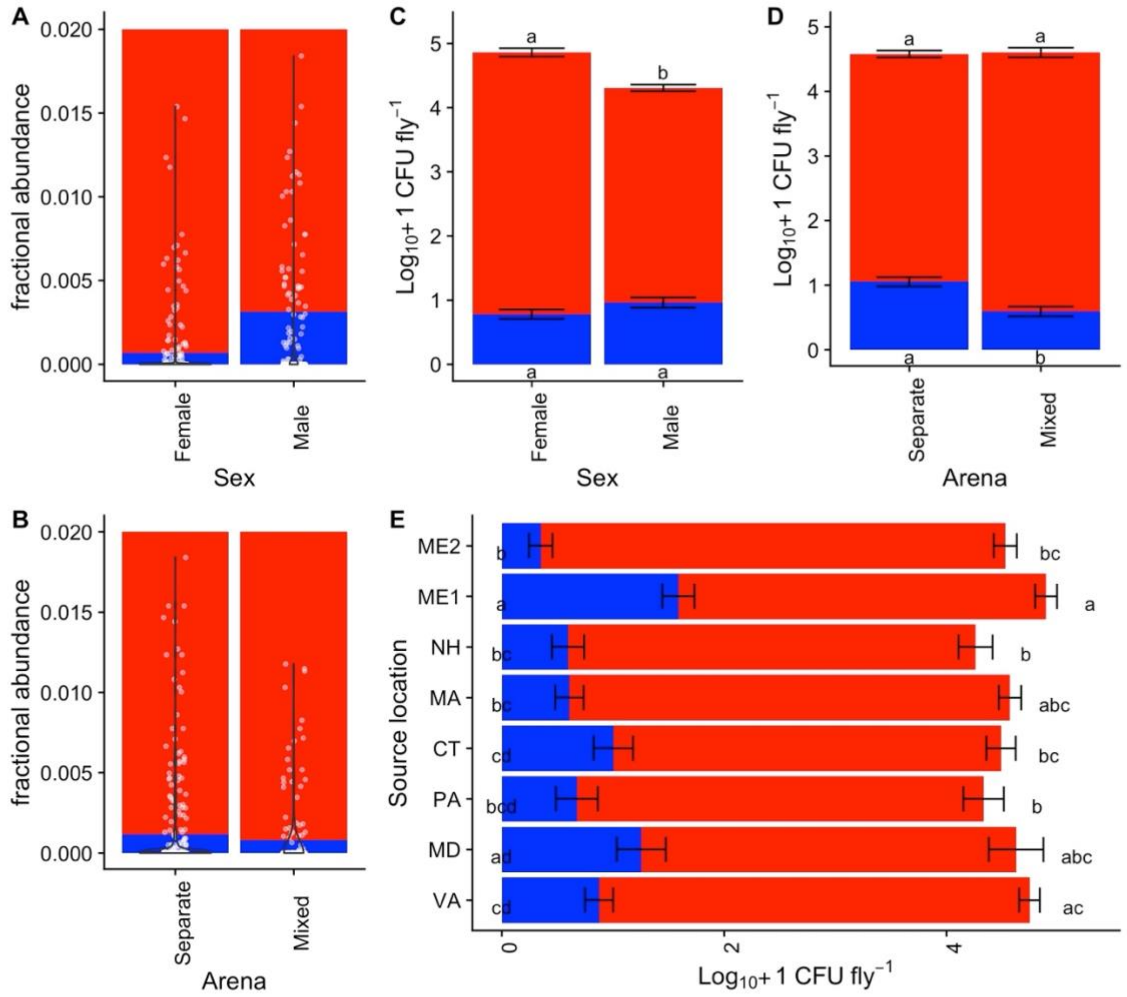


**Figure S2**. The relative abundance of LAB (blue) and AAB (red) in A) male and female CantonS flies inoculated with a 1:1 starting ratio of 3 LAB and 3 AAB strains in the B) separate- or mixed-bacteria condition after three days of transfer to the diet condition plates. We also show the absolute abundance of AAB and LAB in the same flies, separated by C) fly sex, D) diet condition, or E) geography.

Data are shown as the fraction of sample means with overlaid violin plots and the fraction of LAB in each sample as a white dot (A-B) or the mean abundance across all replicates in each of three separate experiments, plus or minus the standard error of the mean (C-E). Compact letter displays next to the red or blue bars report significant differences in the abundance of the AAB or LAB, respectively, between fly populations as determined by a Wilcoxon test (below) with a post-hoc Dunn test. Source locations of the flies are as shown in Figure 1.

Panel C: AAB: $\chi^2_{1, 464} = 41.63$, p = 0; LAB: $\chi^2_{1, 464} = 3.06$, p = 0.08
Panel D: AAB: $\chi^2_{1, 464} = 1.42$, p = 0.2336; LAB: $\chi^2_{1, 464} = 17.78$, p = 0
Panel E: AAB: $\chi^2_{7, 464} = 20.05$, p = 0.01; LAB: $\chi^2_{7, 464} = 51.72$, p = 0

**Table S13. PERMANOVA results for CFU counts (AAB and LAB) of the fly microbiota in a diet-condition experiment with AAB and LAB mixed at a 1:1 ratio before inoculating to diets.**

| | ***Absolute*** | | | | | | ***Relative*** | | | | | |
|---|---|---|---|---|---|---|---|---|---|---|---|---|
| | **Df[a]** | **SS[b]** | **$R^2$** | **F** | **p[c]** | | **Df[a]** | **SS[b]** | **$R^2$** | **F** | **p[c]** | |
| *Arena* | 1.07 | 0.01 | 4.98 | 0.00 | 1.07 | ** | 1 | 0.00 | 0.01 | 3.26 | 0.00 | . |
| *Geography* | 5.29 | 0.04 | 3.51 | 0.00 | 5.29 | *** | 7 | 0.00 | 0.03 | 1.62 | 0.00 | |
| *Sex* | 5.90 | 0.05 | 27.42 | 0.00 | 5.90 | *** | 1 | 0.00 | 0.01 | 4.50 | 0.00 | * |
| *Exp[d]* | 8.64 | 0.07 | 40.17 | 0.00 | 8.64 | *** | 1 | 0.00 | 0.00 | 1.68 | 0.00 | |
| *A * G[e]* | 2.40 | 0.02 | 1.59 | 0.05 | 2.40 | * | 7 | 0.00 | 0.02 | 1.18 | 0.00 | |
| *A * S* | 0.45 | 0.00 | 2.11 | 0.09 | 0.45 | . | 1 | 0.00 | 0.01 | 2.07 | 0.00 | |
| *G * S* | 2.66 | 0.02 | 1.77 | 0.02 | 2.66 | * | 7 | 0.01 | 0.06 | 3.11 | 0.01 | * |
| *Exp / Plate[f]* | 2.77 | 0.02 | 3.22 | 0.00 | 2.77 | *** | 4 | 0.00 | 0.01 | 0.58 | 0.00 | |
| *A * G * S* | 1.67 | 0.01 | 1.11 | 0.32 | 1.67 | | 6 | 0.01 | 0.04 | 2.85 | 0.01 | * |
| *Exp / Plate / Vial[g]* | 24.30 | 0.19 | 1.31 | 0.00 | 24.30 | *** | 80 | 0.03 | 0.23 | 1.15 | 0.03 | |
| *Residual* | 73.31 | 0.57 | | | 73.31 | | 228 | 0.08 | 0.58 | | 0.08 | |

[a] degrees of freedom
[b] sum of squares
[c] p-value
[d] One of two separate experiments in time
[e] "*" = the interaction term
[f] "/" = the nesting term;
[g] vial = the source vial the flies were grown in (30-50 mixed sex flies per vial)

We tested how the dietary microbiota changed in the absence of flies (after inoculating with bacteria for 24h, then culturing at 30°C for 24 h) at two bacterial dilutions (1:1 AAB:LAB, 1:50 AAB:LAB). The rarefied microbiota composition of the diets supplemented with 50X more LAB than AAB was significantly different from the rarefied microbiota composition of the 1:1 ratio inoculated diets ($F_{1, 63}$ = 10.15, $R^2$ = 0.14, $p < 0.003$), and had a higher average of LAB (Fig. S3, Table S14). However, there was not a significant difference in the microbiota composition between arena types (separate- vs. mixed-bacteria condition; $F_{1, 63}$ = 1.11, $R^2$ = 0.02, p = 0.35, the interaction of arena type with LAB concentration was also not significant, $F_{1, 63}$ = 1.51, $R^2$ = 0.02, p = 0.31). The latter finding suggests that the difference between the microbiota of conventional flies introduced to these diets might not be a reflection of the diet; however, this conclusion should be drawn with caution because the fly-free diet analysis reports the microbiota of the diet when the flies are first exposed to it, and the fly microbiota was measured at the end of three consecutive 24-hour periods of 25°C incubation with the diet plates. Regardless, we proceeded to use a 50-fold higher LAB inoculum in our experiments (main text, Figure 2) to help increase the levels of LAB in the flies during the experiments.

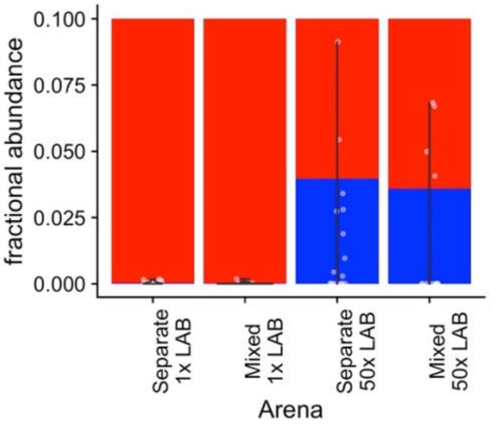


**Figure S3**. The difference in dietary LAB abundance when diet plates are inoculated with 50-fold more LAB than AAB. Data are from four separate experiments and are shown as the fraction of sample means with overlaid violin plots and the fraction of LAB in each sample as a white dot.

The first two columns are from diet plates inoculated with a 1:1 ratio of AAB:LAB. The second two columns are from diet plates inoculated with 50X more LAB than AAB at the start of the experiment.

Diet samples were collected after bacteria were cultured on the plates for 24 h. Diets were incubated without flies in these experiments.

**Table S14. PERMANOVA results for CFU counts (AAB and LAB) of the fly microbiota in a diet-condition experiment with AAB and LAB mixed at a 1:50 ratio before inoculating to diets.**

| | *Absolute* | | | | | | *Relative* | | | | | |
|---|---|---|---|---|---|---|---|---|---|---|---|---|
| | Df[a] | SS[b] | $R^2$ | F | p[c] | | Df[a] | SS[b] | $R^2$ | F | p[c] | |
| *Arena* | 1 | 0.6714 | 0.06 | 12.77 | 0.00 | *** | 1 | 0.00248 | 0.02 | 1.11 | 0.35 | |
| *LAB amount* | 1 | 2.4818 | 0.23 | 47.22 | 0.00 | *** | 1 | 0.022591 | 0.14 | 10.15 | 0.00 | ** |
| *Exp* | 3 | 4.5123 | 0.42 | 28.62 | 0.00 | *** | 3 | 0.010112 | 0.06 | 1.51 | 0.20 | |
| *A * L* | 1 | 0.038 | 0.00 | 0.72 | 0.47 | | 1 | 0.002474 | 0.02 | 1.11 | 0.31 | |
| *Residual* | 57 | 2.9957 | 0.28 | | | | 57 | 0.12691 | 0.77 | | | |
| *Total* | 63 | 10.6991 | 1.00 | | | | 63 | 0.164567 | 1.00 | | | |

**Text S2.** Figures and PERMANOVA tables for each figure shown in the main text, except for absolute CFU counts in the flies.

*Corresponds to Figure 2B and Table S1*

**Table S15**. **PERMANOVA results for CFU counts (AAB and LAB) of gnotobiotic flies in eight populations originally collected from each of eight different locations in the eastern USA**. Data are shown as the mean abundance of AAB (red) and LAB (blue) across all replicates in each of three separate experiments, plus or minus the standard error of the mean. Compact letter displays next to the red or blue bars report significant differences in the abundance of the AAB or LAB, respectively, between fly populations as determined by a Kruskal-Wallis test with post-hoc multiple comparisons by a Dunn test (AAB: $\chi^2_{7, 225} = 10.33$, p = 0.17; LAB: $\chi^2_{7, 225} = 20.39$, p = 0.005). Source locations of the flies are as shown in Figure 1.

| | *df* [a] | *SS* [b] | *R²* | *F* | *p* [c] | |
|---|---|---|---|---|---|---|
| *Geography* | 7 | 3.50 | 0.07 | 2.59 | 0.001 | *** |
| *Sex* | 1 | 0.49 | 0.01 | 2.56 | 0.04 | * |
| *Exp* [d] | 2 | 1.23 | 0.02 | 3.18 | 0.01 | * |
| *Plate* [e] | 7 | 3.40 | 0.07 | 2.52 | 0.002 | ** |
| *G * S* [f] | 7 | 1.21 | 0.02 | 0.90 | 0.56 | |
| *Plate / Vial* [g] | 90 | 20.06 | 0.39 | 1.16 | 0.09 | . |
| *Residual* | 111 | 21.40 | 0.42 | | | |
| *Total* | 225 | 51.28 | 1.00 | | | |

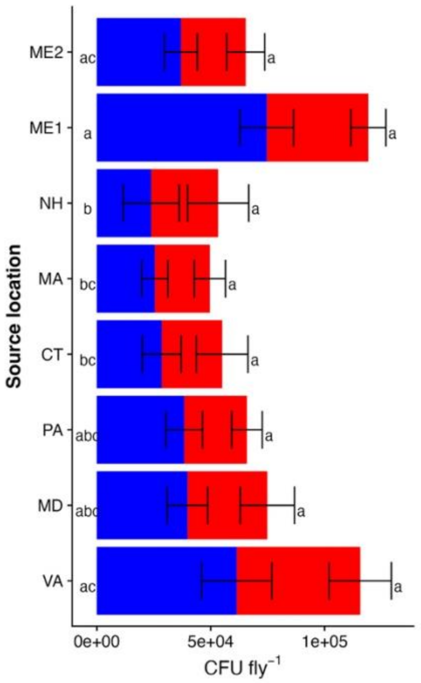


[a] degrees of freedom
[b] sum of squares
[c] p-value
[d] One of three separate experiments in time
[e] 96-well plate on which individual replicates of flies were homogenized and dilution plated;
[f] "*" = the interaction term
[g] the source vial the flies were grown in (30-50 mixed sex flies per vial); "/" = the nesting term

Test statistics:

*Corresponds to Figure 3C and Table S2*

**Table S16. PERMANOVA results for CFU counts (AAB and LAB) of gnotobiotic flies from the eight populations in our study, exposed to mixed- and separated-bacteria diet conditions.** Data are shown as the mean abundance of AAB (red) and LAB (blue) across all replicates in each of two separate experiments, plus or minus the standard error of the mean. A) absolute LAB counts only; B) stacked bar chart of the absolute LAB and AAB counts.Compact letter displays next to the red or blue bars report significant differences in the abundance of the AAB or LAB, respectively, between fly populations as determined by a Kruskal-Wallis test with post-hoc multiple comparisons by a Dunn test (AAB: $\chi^2_{1, 393}$ = 1.1, p = 0.30; LAB: $\chi^2_{1, 393}$ = 156.35, p < $10^{-4}$).

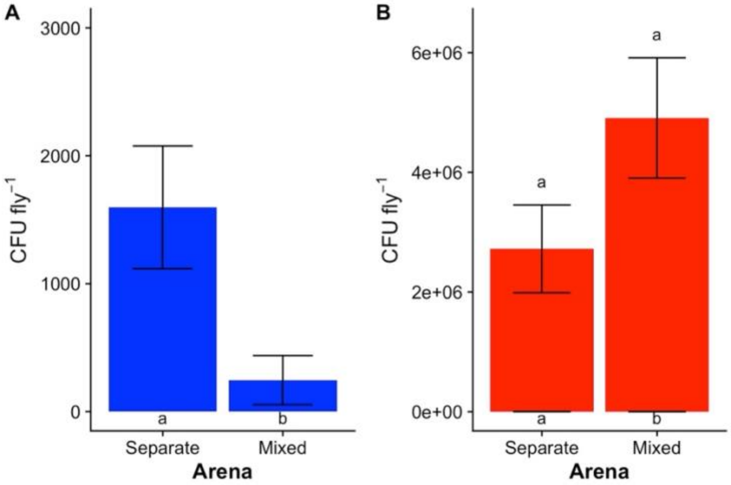


| | *df* [a] | *SS* [b] | *R²* | *F* | *p* [c] | |
|---|---|---|---|---|---|---|
| *Arena* [d] | 1 | 0.36 | 0.00 | 2.35 | 0.066 | . |
| *Sex* | 1 | 1.55 | 0.02 | 10.17 | 0.001 | *** |
| *Geography* | 7 | 3.74 | 0.04 | 3.50 | 0.001 | *** |
| *Exp* [e] | 1 | 4.07 | 0.04 | 26.72 | 0.001 | *** |
| *A * S* [f] | 1 | 0.03 | 0.00 | 0.17 | 0.956 | |
| *E / Plate* [g] | 2 | 21.36 | 0.21 | 70.05 | 0.001 | *** |
| E / P / Arena [h] | 69 | 20.85 | 0.21 | 1.98 | 0.001 | *** |
| Residual | 311 | 47.42 | 0.48 | | | |
| *Total* | 393 | 99.39 | 1.00 | | | |

[a] degrees of freedom
[b] sum of squares
[c] p-value
[d] mixed- or separated-bacteria condition
[e] One of two separate experiments in time
[f] * = the interaction term
[g] 96-well plate on which individual replicates of flies were homogenized and dilution plated; / represents the nesting term
[h] the source arena the flies were collected from

*No tables or figures based on absolute abundances correspond to Figure 3C and Tables S3-4 because the absolute quantity of diet was not scaled at the time of collection and can only be used for relative abundance analysis.*

*Corresponds to Figure 3D and Table S5*

**Table S17. PERMANOVA results for CFU counts (AAB and LAB) of flies in the mixed bacteria arenas.** Data are shown as the log10-transformed mean abundance of AAB (red) and LAB (blue) across all replicates in each of two separate experiments, plus or minus the log10-transformed standard error of the mean. Compact letter displays next to the red or blue bars report significant differences in the abundance of the AAB or LAB, respectively, between fly populations as determined by a Kruskal-Wallis test with post-hoc multiple comparisons by a Dunn test (AAB: $\chi^2_{7, 197} = 33.01$, $p < 10^{-4}$; LAB: $\chi^2_{7, 197} = 26.68$, $p = 0.0002$).

| | df [a] | SS [b] | $R^2$ | F | p [c] | |
|---|---|---|---|---|---|---|
| *Geography* | 7 | 4.78 | 0.09 | 4.49 | 0.001 | *** |
| *Sex* | 1 | 0.73 | 0.01 | 4.82 | 0.002 | ** |
| *Exp* [d] | 1 | 2.06 | 0.04 | 13.51 | 0.001 | *** |
| *G * S* [e] | 7 | 0.90 | 0.02 | 0.85 | 0.66 | |
| *E / Plate* [f] | 3 | 13.44 | 0.26 | 29.44 | 0.001 | *** |
| *E / P / Arena* [g] | 28 | 6.78 | 0.13 | 1.59 | 0.001 | *** |
| *Residual* | 150 | 22.83 | 0.44 | | | |
| *Total* | 197 | 51.53 | 1.00 | | | |

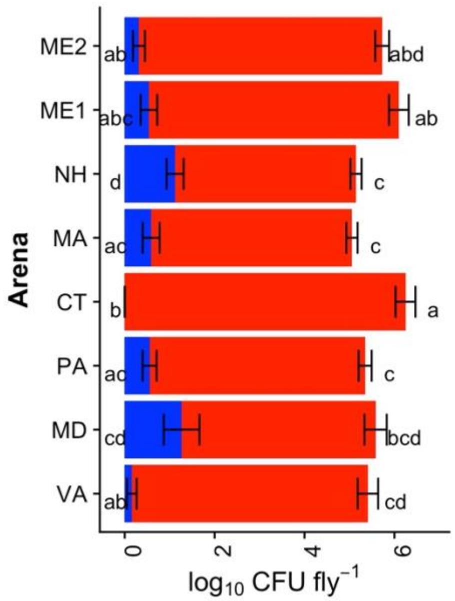


[a] degrees of freedom
[b] sum of squares
[c] p-value
[d] One of two separate experiments in time
[e] * = the interaction term
[f] plate = 96-well plate on which individual replicates of flies were homogenized and dilution plated; / represents the nesting term
[g] Arena = the source arena flies were collected from, one of three arenas per condition in each experiment

*Corresponds to Figure 3E and Table S6*

**Table S18. PERMANOVA results for CFU counts (AAB and LAB) of flies in the separated bacteria arenas.** Data are shown as the log10-transformed mean abundance of AAB (red) and LAB (blue) across all replicates in each of two separate experiments, plus or minus the log10-transformed standard error of the mean. Compact letter displays next to the red or blue bars report significant differences in the abundance of the AAB or LAB, respectively, between fly populations as determined by a Kruskal-Wallis test with post-hoc multiple comparisons by a Dunn test (AAB: $\chi^2_{7, 195} = 33.75$, $p < 10^{-4}$; LAB: $\chi^2_{7, 195} = 15.79$, $p = 0.0271$).

| | *df* [a] | *SS* [b] | $R^2$ | *F* | *p* [c] | |
|---|---|---|---|---|---|---|
| *Geography* | 7 | 4.70 | 0.10 | 4.31 | 0.001 | *** |
| *Sex* | 1 | 0.80 | 0.02 | 5.15 | 0.002 | ** |
| *Exp* [d] | 1 | 2.45 | 0.05 | 15.71 | 0.001 | *** |
| *G * S* [e] | 7 | 0.96 | 0.02 | 0.88 | 0.62 | |
| *E / Plate* [f] | 3 | 9.78 | 0.21 | 20.92 | 0.001 | *** |
| *E / P / Vial* [g] | 29 | 5.90 | 0.12 | 1.30 | 0.04 | * |
| *Residual* | 147 | 22.91 | 0.48 | | | |
| *Total* | 195 | 47.50 | 1.00 | | | |

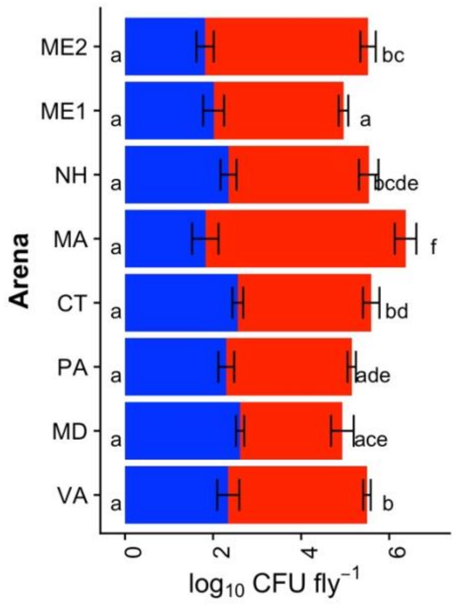


[a] degrees of freedom
[b] sum of squares
[c] p-value
[d] mixed- or separated-bacteria condition
[e] flies or diets
[f] One of two separate experiments in time
[g] * = the interaction term
[h] plate = 96-well plate on which individual replicates of flies were homogenized and dilution plated; / represents the nesting term

*Corresponds to Figure 4A and Table S7*

**Table S19. PERMANOVA results for CFU counts (AAB and LAB) of the total and resident-only portions of gnotobiotic flies from the eight populations in our study.** Data are shown as the mean abundance of AAB (red) and LAB (blue) across all replicates in each of three separate experiments, plus or minus the standard error of the mean. Compact letter displays next to the red or blue bars report significant differences in the abundance of the AAB or LAB, respectively, between fly populations as determined by a Kruskal-Wallis test with post-hoc multiple comparisons by a Dunn test (AAB: $\chi^2_{1, 426} = 28.77$, $p < 10^{-4}$; LAB: $\chi^2_{1, 426} = 20.69$, $p < 10^{-4}$).

| | *df* [a] | *SS* [b] | $R^2$ | *F* | *p* [c] | |
|---|---|---|---|---|---|---|
| Geography | 7 | 2.58 | 0.03 | 1.96 | 0.005 | ** |
| Sex | 1 | 0.36 | 0.00 | 1.92 | 0.104 | |
| Niche [d] | 1 | 2.61 | 0.03 | 13.90 | 0.001 | *** |
| Exp [e] | 2 | 1.36 | 0.01 | 3.62 | 0.002 | ** |
| Plate [f] | 15 | 8.47 | 0.08 | 3.01 | 0.001 | *** |
| G * S [g] | 7 | 1.29 | 0.01 | 0.98 | 0.489 | |
| G * N | 7 | 2.24 | 0.02 | 1.71 | 0.034 | * |
| S * N | 1 | 0.26 | 0.00 | 1.37 | 0.235 | |
| Plate / Vial [h] | 183 | 45.60 | 0.44 | 1.33 | 0.001 | *** |
| Residual | 202 | 37.92 | 0.37 | | | |
| *Total* | 426 | 102.69 | 1.00 | | | |

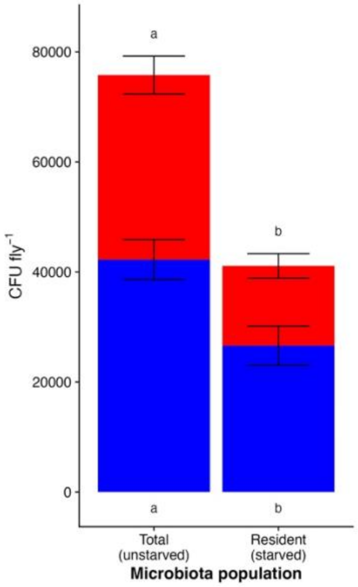


[a] degrees of freedom
[b] sum of squares
[c] p-value
[d] flies in the directly-homogenized (total microbes) or briefly-starved (only resident microbes) treatments
[e] One of three separate experiments in time
[f] 96-well plate on which individual replicates of flies were homogenized and dilution plated
[g] "*" = the interaction term
[h] "/" = the nesting term; the source vial the flies were grown in (30-50 mixed sex flies per vial)

*Corresponds to Figure 4B and Table S8*

**Table S20. PERMANOVA results for CFU counts (AAB and LAB) of the resident-only portion of gnotobiotic flies from the eight populations in our study.** Data are shown as the mean abundance of AAB (red) and LAB (blue) across all replicates in each of three separate experiments, plus or minus the standard error of the mean. Compact letter displays next to the red or blue bars report significant differences in the abundance of the AAB or LAB, respectively, between fly populations as determined by a Kruskal-Wallis test with post-hoc multiple comparisons by a Dunn test (AAB: $\chi^2_{7,\ 200} = 8.91$, p = 0.26; LAB: $\chi^2_{7, 200} = 8.87$, p = 0.26).

| | *df* [a] | *SS* [b] | $R^2$ | *F* | *p* [c] | |
|---|---|---|---|---|---|---|
| Geography | 7 | 1.66 | 0.03 | 1.31 | 0.13 | |
| Sex | 1 | 0.14 | 0.00 | 0.75 | 0.51 | |
| Exp [d] | 2 | 1.33 | 0.03 | 3.67 | 0.00 | *** |
| Plate[e] | 6 | 3.84 | 0.08 | 3.52 | 0.00 | *** |
| G * S [f] | 7 | 1.76 | 0.04 | 1.38 | 0.11 | |
| Plate / Vial [g] | 86 | 23.8 | 0.49 | 1.53 | 0.00 | *** |
| Residual | 91 | 16.5 | 0.34 | | | |
| *Total* | 200 | 49.0 | 1.00 | | | |

[a] degrees of freedom
[b] sum of squares
[c] p-value
[d] One of three separate experiments in time
[e] 96-well plate on which individual replicates of flies were homogenized and dilution plated
[f] "*" = the interaction term
[g] "/" = the nesting term; the source vial the flies were grown in (30-50 mixed sex flies per vial)

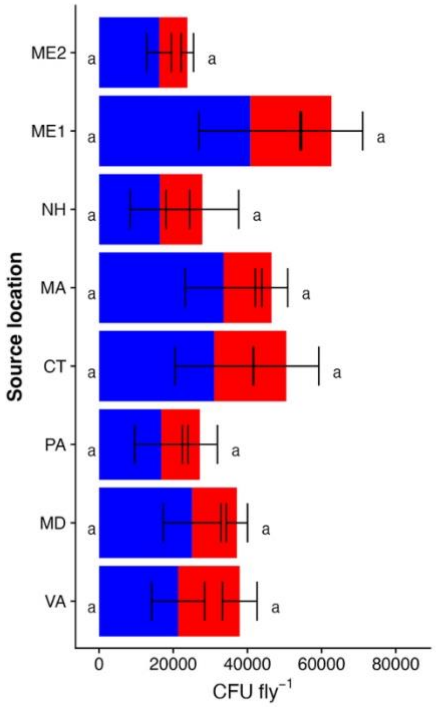

*Corresponds to Figure 4C*

**FigureS1_TextS2.** Data are shown as the mean abundance of AAB (red) and LAB (blue) across all replicates in each of three separate experiments, plus or minus the standard error of the mean. Compact letter displays next to the red or blue bars report significant differences in the abundance of the AAB or LAB, respectively, between fly populations as determined by a Kruskal-Wallis test with post-hoc multiple comparisons by a Dunn test (AAB: $\chi^2_{7,\ 26} = 5.49$, $p = 0.60$; LAB: $\chi^2_{7,27} = 13.72$, $p = 0.056$).

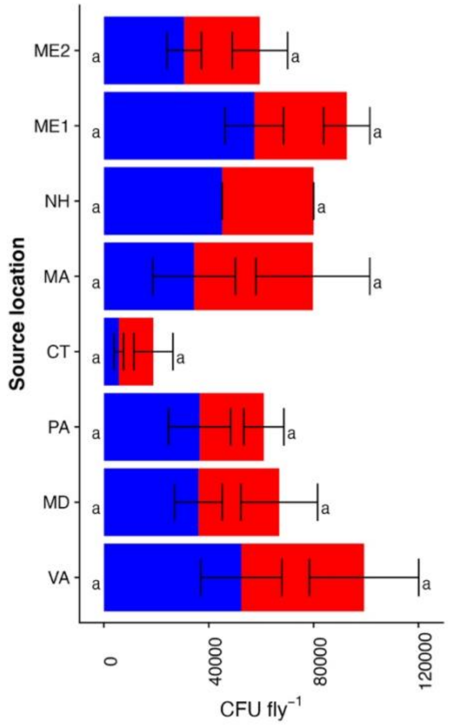

*Corresponds to Figure 5A and Table S9*

**Table S21. PERMANOVA results for CFU counts (AAB and LAB) of the total microbiota in gnotobiotic flies from the eight populations in our study when reared at 50% and 60-75% RH.** Data are shown as the mean abundance of AAB (red) and LAB (blue) across all replicates in each of three separate experiments, plus or minus the standard error of the mean. Compact letter displays next to the red or blue bars report significant differences in the abundance of the AAB or LAB, respectively, between fly populations as determined by a Kruskal-Wallis test with post-hoc multiple comparisons by a Dunn test (AAB: $\chi^2_{7,\ 473} = 23.04$, $p < 10^{-4}$; LAB: $\chi^2_{7,\ 473} = 23.56$, $p = 0.001$).

| | *df* [a] | *SS* [b] | *R²* | *F* | *p* [c] | |
|---|---|---|---|---|---|---|
| *Geography* | 7 | 3.91 | 0.04 | 3.34 | 0.00 | *** |
| *Sex* | 1 | 0.33 | 0.00 | 1.99 | 0.09 | . |
| *Humidity* [d] | 1 | 0.83 | 0.01 | 4.97 | 0.00 | ** |
| *Exp* [e] | 2 | 1.14 | 0.01 | 3.41 | 0.00 | ** |
| *Plate* [f] | 9 | 6.13 | 0.06 | 4.07 | 0.00 | *** |
| *G * S* [g] | 7 | 0.80 | 0.01 | 0.68 | 0.85 | |
| *G * H* | 7 | 2.79 | 0.03 | 2.38 | 0.00 | ** |
| *S * H* | 1 | 0.14 | 0.00 | 0.83 | 0.45 | |
| *P / Vial* [h] | 188 | 41.5 | 0.42 | 1.32 | 0.00 | ** |
| *Residual* | 250 | 41.7 | 0.42 | | | |
| *Total* | 473 | 99.4 | 1.00 | | | |

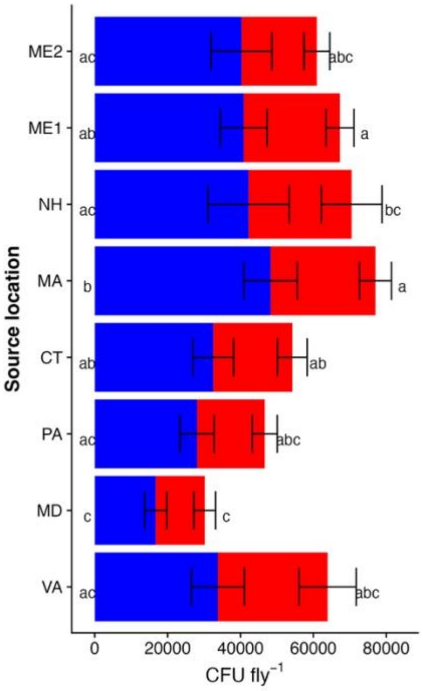


[a] degrees of freedom
[b] sum of squares
[c] p-value
[d] flies reared at ambient (~25%) RH, at 50% RH, or at 60-75% RH
[e] One of three separate experiments in time
[f] 96-well plate on which individual replicates of flies were homogenized and dilution plated
[g] "*" = the interaction term
[h] "/" = the nesting term; vial = the source vial the flies were grown in (30-50 mixed sex flies per vial)

*Corresponds to Figure 5B and Table S10*

**Table S22. PERMANOVA results for CFU counts (AAB and LAB) of the resident microbiota in gnotobiotic flies from the eight populations in our study when reared at 50% and 60-75% RH.** Data are shown as the mean abundance of AAB (red) and LAB (blue) across all replicates in each of three separate experiments, plus or minus the standard error of the mean. Compact letter displays next to the red or blue bars report significant differences in the abundance of the AAB or LAB, respectively, between fly populations as determined by a Kruskal-Wallis test with post-hoc multiple comparisons by a Dunn test (AAB: $\chi^2_{7,\ 440}$ = 5.63, p = 0.58; LAB: $\chi^2_{7,\ 440}$ = 5.4, p = 0.61).

| | *df* [a] | *SS* [b] | $R^2$ | *F* | *p* [c] | |
|---|---|---|---|---|---|---|
| *Geography* | 7 | 2.07 | 0.02 | 1.38 | 0.10 | |
| *Sex* | 1 | 0.45 | 0.00 | 2.12 | 0.08 | . |
| *Humidity* | 1 | 0.39 | 0.00 | 1.82 | 0.13 | |
| *Exp* | 2 | 1.61 | 0.01 | 3.77 | 0.00 | ** |
| *Plate* | 8 | 6.34 | 0.06 | 3.71 | 0.00 | *** |
| *G * S* | 7 | 1.97 | 0.02 | 1.32 | 0.15 | |
| *G * H* | 7 | 3.50 | 0.03 | 2.34 | 0.00 | *** |
| *S * H* | 1 | 0.22 | 0.00 | 1.04 | 0.34 | |
| *Plate / Vial* | 185 | 47.0 | 0.42 | 1.19 | 0.01 | * |
| *Residual* | 221 | 47.1 | 0.43 | | | |
| *Total* | 440 | 110. | 1.00 | | | |

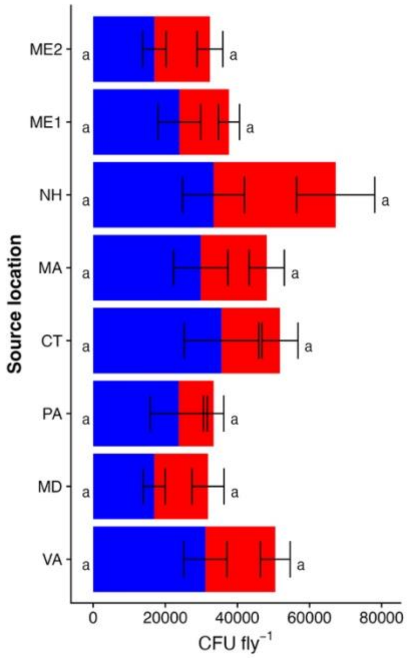


[a] degrees of freedom
[b] sum of squares
[c] p-value
[d] flies reared at ambient (~25%) RH, at 50% RH, or at 60-75% RH
[e] One of three separate experiments in time
[f] 96-well plate on which individual replicates of flies were homogenized and dilution plated
[g] "*" = the interaction term
[h] "/" = the nesting term; vial = the source vial the flies were grown in (30-50 mixed sex flies per vial)

*Corresponds to Figure 5C*

**FigureS2_TextS2.** Data are shown as the mean abundance of AAB (red) and LAB (blue) across all replicates in each of three separate experiments, plus or minus the standard error of the mean. Compact letter displays next to the red or blue bars report significant differences in the abundance of the AAB or LAB, respectively, between fly populations as determined by a Kruskal-Wallis test with post-hoc multiple comparisons by a Dunn test (AAB: $\chi^2_{7,\ 50} = 6.7$, p = 46; LAB: $\chi^2_{7, 50} = 12.71$, p = 0.08).

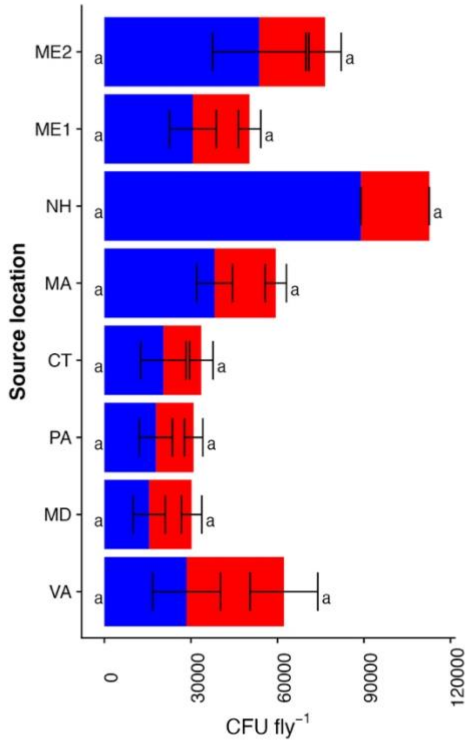
